# Simulated climate change magnifies genetic vulnerabilities from mutation load and maladaptation

**DOI:** 10.1101/2025.09.25.677362

**Authors:** Laura Leventhal, Megan R. Ruffley, Carnegie Field Consortium, Moises Exposito-Alonso

## Abstract

As climate change intensifies, the genetic diversity and composition of natural populations will become critical for adaptation and survival. Standing genetic diversity within populations differs across a species’ range, due to past demographic and natural selection processes driving the accumulation of adaptive, neutral, and deleterious variation. While accumulating genomic knowledge could be used to evaluate population extinction risk from local mal-adaptive genetic makeups, testing such approaches in natural environments remains challenging. Leveraging the genomic resources of *Arabidopsis thaliana*, we created experimental synthetic populations of similar genetic diversity but differing genetic makeups by mixing 245 natural accessions with different levels of potentially-climate-adaptive alleles and/or genomic burden of deleterious mutations. We planted our populations in a climate change field experiment simulating a gradient of declining rainfall. By tracking survival and reproduction of 135 synthetic experimental populations over three years, we show substantial predictability of genetic makeup on survival and population growth rate. Further, the accumulation of deleterious mutations and locally (mal)adaptive alleles synergistically reduces fitness in increasingly stressful climates. Our findings underscore that for populations to have the greatest chance of surviving climate change, the optimal combination of genomic makeups is essential.

## Introduction

Species face increasingly variable climates to which they must adapt to prevent extinction ^1,2^. Standing genetic variation within populations fuels adaptation ^3,4^, but natural populations within a species’ range often not only vary in overall number and diversity of genetic variants but also the composition of adaptive or deleterious alleles—i.e. genetic makeup^5–7^. Such differences in genetic makeup can be shaped by a combination of past natural selection, demographic history, and admixture, and can determine how populations respond to global environmental change ^8–11^. Divergence in genetic makeup across populations arises through local adaptation across a species’ native range ^12,13^, as populations inhabiting different environments progressively accumulate alleles beneficial precisely in their local environments. However, past demographic processes also cause divergence in wild populations including accumulation of deleterious mutations that may decrease fitness, such as through population isolation and inbreeding ^14,15^. Ultimately, the combination of such globally deleterious and locally adaptive genetic variation will determine the readiness of populations to adapt to future conditions.

In plant species such as the model system *Arabidopsis thaliana—*where differences in geographic and evolutionary patterns are pronounced—large-scale genomic efforts are beginning to reveal the distribution of both genetic diversity and makeup across broad landscapes ^16–21^. For instance, genomic screens of local adaptation have revealed that range edge populations often contain more adaptive variants under extreme conditions such as drought ^18^, consistent with ecological predictions across species that lower latitude populations should carry greater adaptive potential^8,17,22–26^. This is in contrast to the alternate hypothesis that isolation of populations in edge environments could also lead to demographic processes that reduce the efficiency of selection. Small populations experience stronger drift relative to purifying selection, leading to an increase in the burden of deleterious alleles and a reduction in mean fitness (i.e., genetic load) ^27^, which can be captured by counting genome-wide nonsynonymous mutations such as stop codons and amino-acid relative to silent synonymous mutations (*P_n_/P_s_* ^28,29^). In *A. thaliana*, it appears that there is a gradient of increasing mutations towards post-glacial northern edges ^30^, a pattern following polar trends observed in other plant species including *A. lyrata, Mercurialis annua, Solanum chilense,* and *Vitis arizonica* ^29,31–33^.

Although the roles of adaptive variants and local adaptation in fitness are well established ^27,34–36^, the combined effects of local adaptation and mutation load—particularly in the face of rapid environmental change—remain almost entirely unexplored ^37,38^ and thus impossible to forecast. On one hand, rapid climate change is making genetic variants that were once advantageous become less well-suited to the new environment, which is attempted to be captured by new genomic prediction tools of genomic offset and polygenic scores ^39^. While on the other hand, inbreeding and deleterious load is of utmost conservation concern for at-risk populations and is often measured utilizing mutation annotation tools ^34,40^. While it could be hypothesized that mutation load and locally (mal)adaptive alleles may amplify, counteract, or interact in complex ways (additive, synergistic, or antagonistic) ^41^, the interactions between these factors remain poorly quantified. Ultimately, a comprehensive understanding of these two dimensions of genetic makeup is essential in making predictions of fitness in the future complex climate change landscape ^42^ but it has remained elusive given both genomic and experimental limitations.

To disentangle how the two dimensions of genetic composition including local maladaptation and genetic load affect a species’ response to changing climates, we designed experimental synthetic populations of *A. thaliana* using several genomic prediction methods and tested their survival under complex climate change precipitation manipulations. Because precipitation trajectories are expected to become highly stochastic with climate change, threatening plant communities ^43–45^, we recreated a fully-factorial set of precipitation regimes. In parallel, we created the experimental synthetic populations leveraging *A. thaliana’s* genomic resources ^46^, selecting groups of accessions with comparable genetic diversity but divergent genomic vulnerability metrics of mutation load ^28^ and polygenic scores of local mal-adaptation ^18,47^. We found that our predictions of local adaptation and mutation load significantly explained survival in a synergistic fashion with increased predictability of individual’s fitness with progressively stressful climate conditions, and that such variation in turn had a ripple effect on population growth rate for up to three generations. These results underscore the importance of the type of genetic diversity across populations to understand their complex responses to climate change.

## Results & Discussion

### Precipitation stochasticity drives mortality in *A. thaliana*

To understand how climate change-like precipitation alterations may impact the survival of a temperate annual plant species, *Arabidopsis thaliana,* we designed a fully factorial rainfall manipulation experiment using an automated irrigation system at the Carnegie common garden in Stanford University (<u>37°25’42.6"N, 122°10’45.8"W</u>, Stanford, California, USA, **Fig. 1A**). The field station represents a Mediterranean climate (average max annual temperature = 21.1°C, min = 8.4°C, annual average precipitation = 394mm from NOAA Palo Alto, CA, US Climate Normals from 1991-2020, **Figure S1**). This climate is most similar to the southern edge of the species‘ range in the Mediterranean region and the Levant. In order to mimic rainfall abundance and variability in *A. thaliana*’s natural range, we programmed twelve of the fourteen treatments to receive rainfall additions, (**Fig. S2-3**) specifically 300, 600, or 1200mm throughout the season. Through the rainfall addition scheme, we varied the frequency of rainfall to monthly, weekly, biweekly, and every other day. The control treatment received the total natural rainfall for the 2021-2022 season (230mm), while the 50% reduction received half the rainfall for the season (**Fig. 1E, S4**), and both of these treatments (the natural and 50% reduction) received natural stochastic rainfall frequencies experienced during the growth season 2021-2022 (**Fig. 1E**). Our precipitation regime successfully mirrored the local precipitation of the accessions’ geographic origin, thus confirming we recapitulated precipitation-driven local adaptation (**Fig. 1D, S6**).

**Figure 1.**
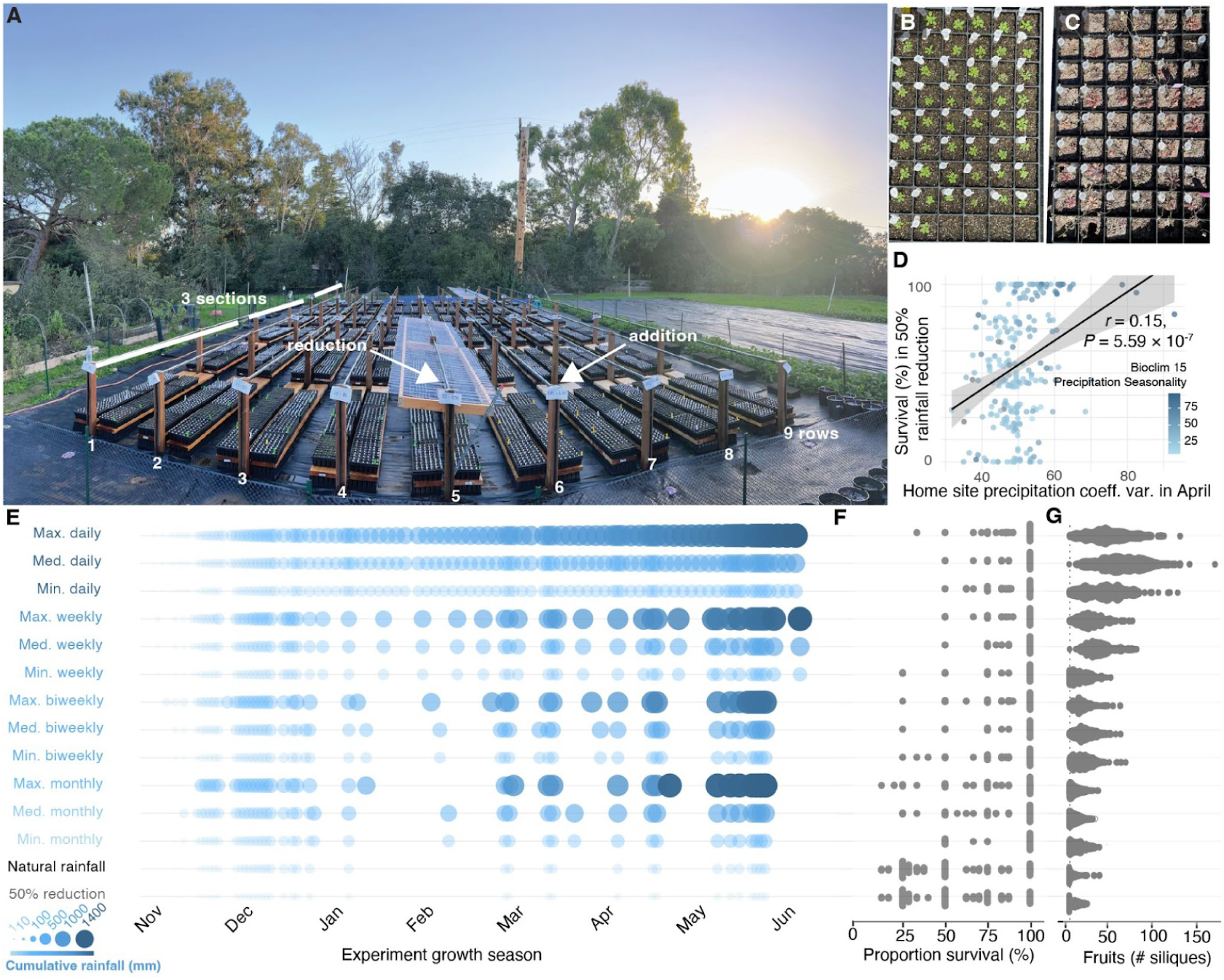
**Simulation of a fully-factorial rainfall manipulation recreates a fine plant mortality gradient.**(**A**) Overview of the experimental site in the Carnegie Institution for Science field site (Stanford, CA, USA). The site had 9 rows with 3 sections per row, and each of the 27 sections was equipped with an independent rainfall irrigation system (addition). Additionally, two 50% rainfall exclusion structures (reduction) were installed. (**B**) Example tray from high rainfall treatment during the rosette stage (tubes originally used to sow the seeds remained as accession identification tags). (**C**) Example tray from precipitation reduction treatment in rosette and flowering stage. (**D**) Relationship between survival in most extreme drought treatment (50% reduction) and coefficient of variation in precipitation from home site in the month of April. Points are colored by Bioclim variable 15 (precipitation seasonality). (**E**) Rainfall addition experimental design for the 14 precipitation levels (visualized cumulatively), including variations in both frequency (distance between circles) and abundance (size of circles) of rainfall addition events. Daily rainfall received water every other day and by the end of the season received the maximum amount of water. Other imposed frequencies were weekly, biweekly, and monthly. Three potential abundances were maximum (1200mm), medium (600mm), and minimum (300mm), distributed temporally given the previous frequency treatments; fully factorially. No-addition, no-exclusion treatment (natural California) received the natural rainfall of the 2021-2022 growing season (231mm), and a 50% reduction treatment received approximately half (115.5mm). (**F**) Distribution of percent survival and (**G**) silique number for all accessions by treatment.

Within each rainfall treatment, we planted in high replication experimental synthetic populations consisting of a mix of accessions of *A. thaliana* with a genetic design informed by genomic prediction metrics but roughly similar level of diversity (**Methods 2**. Accession selection, **Table S1**). We planted synthetic populations in trays with 60 cells with one plant per cell, thus reducing direct space competition at the experiment’s start (**Fig. 1B-C**). These synthetic population trays can be thought of as a small patch or herb population that may or may not survive the imposed environmental stress. We then tracked whether individual plants (n=14,240) survived and counted their fruit set. Survival was sharply different across precipitation treatments, (X^2^ = 2248.7, df = 13, *P* < 2.2 × 10^-16^) with the most water abundant environments naturally permitting high survival (89% survival on average [80-94%]), while water-limited treatments roughly killed half the individuals across synthetic populations (53% survival [25%-83%]) (**Fig. 1F**). Fruit set was also variable across rainfall treatments (Kruskal-Wallis H = 429.5, p < 2 × 10^-16^, **Fig. 1G, Table S2**). Making use of the full-factorial combinations of abundance and stochasticity of rainfall, we remarkably found that total rainfall during the growth season had little effect on either survival or seed set (proportion of variance explained [PVE] = 3.2%). Instead, frequency of rainfall explained most variance in survival (PVE = 46.2%) (**Table S3**), in agreement with ecological understanding that frequency of droughts drives mortality and adaptation in annual plants ^48–53^. In addition, fitness decayed non-linearly over the precipitation gradient (y = -0.075 -110x - 58x^2^, p = 2× 10^-16^, marginal R^2^ = 0.19, **Fig. 1F**); that is, the simulated worsening climate conditions may create a tipping point of mortality, which has been theorized to drive the limits of adaptation at species range edges ^54–56^. Similarly, the relationship between silique number and treatment ranked by stress was significantly quadratic (y = 0.020 -79x + 1.1x^2^, p = 2× 10^-16^, marginal R^2^ = 0.77, **Fig. 1G**). While we observed a correlation between survival and accessions’ climate of origin (**Fig. 1D, S6**), it is still unclear how much genetic makeup could cultivate such a mortality tipping point, and whether genomics could enable the predictions of population vulnerability with climate change ^57^.

### Predictions of (mal)adaptive genetic makeup indicate substantial fitness declines under increasingly severe rainfall conditions

To understand how differences in the genetic makeup of populations may increase the risk of local extinction, we studied several genomic (mal)adaptation dimensions using *A. thaliana*’s genetic resources. First, we used Genome-Wide Associations (GWA) of fitness traits in a previous rainfall reduction common garden (Madrid, Spain) ^18^, which permitted polygenic score predictions of survival under range edge conditions (*α_score_ = Σg_i_s_i_*, where *α_score_* represents the weighted sum of the effect of genetic variants, *g_i_*is the genotype of variant *i* and *s_i_* is the estimated effect size of that variant on survival, summing across all relevant loci, see **Methods**). This “adaptation score”, showed a significant quadratic relationship with the accessions’ latitude of origin after relatedness control, which confirms previous reports ^58^ indicating accessions at both North and South edges may withstand more extreme environments even when correcting for population structure (y = 8.33× 10^-7^ + 0.00271x + 0.01872x^2^, p_quadratic_ = 2.2 × 10^-16^, marginal R^2^ = 0.54, **Fig. 2A**, see discussion on population structure **Text S1**). Comparing fitness of experimental populations with a mix of high versus low adaptation score genotypes, we corroborated that synthetic experimental populations with high adaptation score had significantly increased fitness in eight of the fourteen treatments on the dry end of the distribution (**Fig. 2B**), with the strongest difference in 50% reduction rainfall condition (survival _high α_= 78% [CI95% = 71% - 84%], s _low α_ = 38% [CI95% = 30% - 45%], Wilcoxon 3.78 × 10^4^, p = 1.59 × 10^-18^, **Table S4**) and no observable trade-off (**Fig. 2B**).

**Figure 2.**
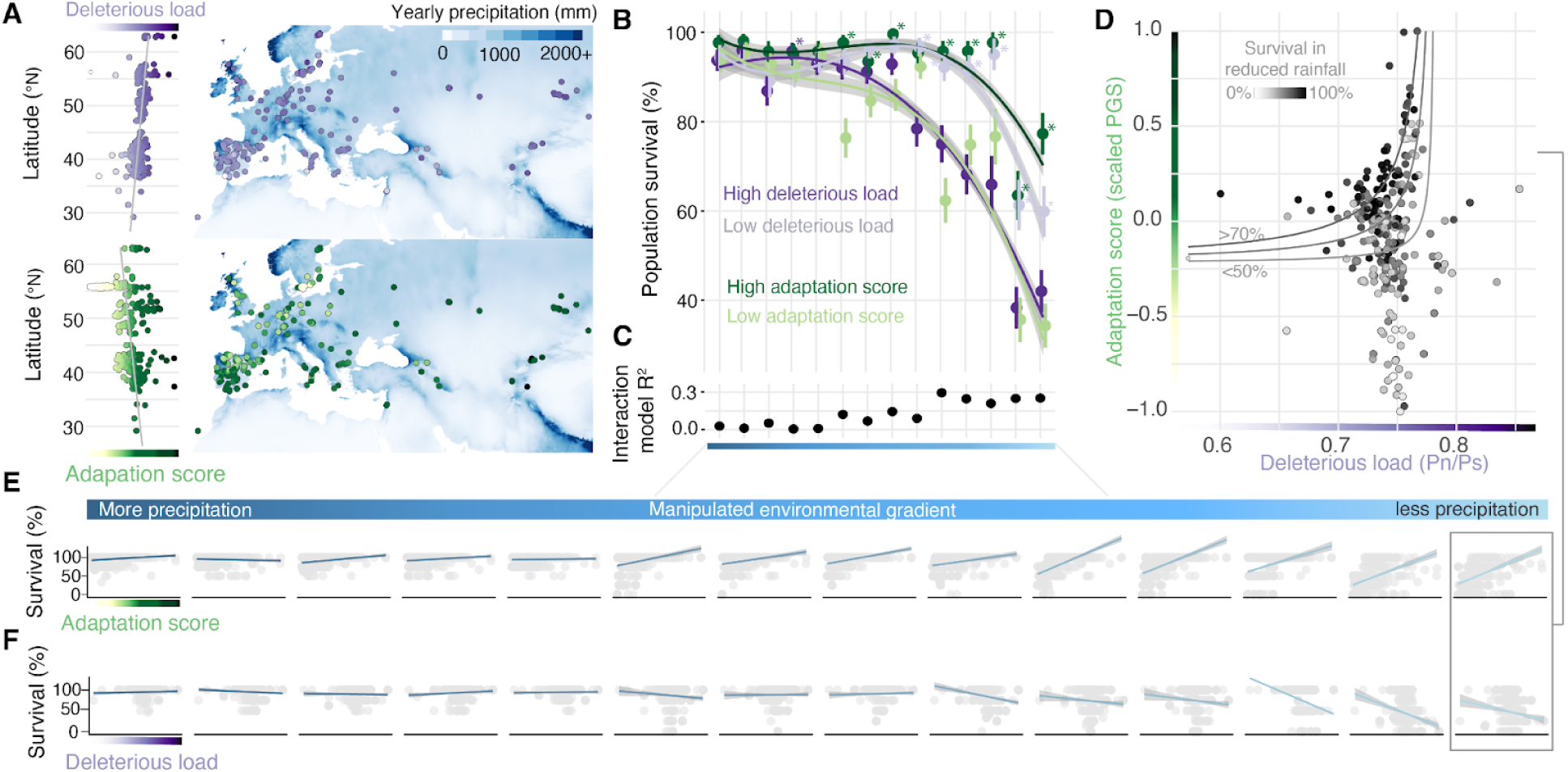
Mutation load and polygenic local adaptation scores correlate and interact with fitness across a climate gradient. (**A**) Maps of all accessions from the experiment with points colored by mutation load and adaptation score by latitude and across the geographic range. Annual precipitation (mm) across Eurasia and North Africa is shown by a blue color gradient. (**B**) Differences in population percentage survival by treatment. Grouped colors represent the high and low extremes of the paired populations (high and low genetic load (purples) and high and low adaptation score (green). Asterisks (*) of the same color family as the population indicate that the Wilcoxon rank difference between the two paired colors (greens, purples) are significantly different, and the asterisk is placed next to the significantly higher fitness population. **(C)** Proportion of variance explained (R^2^) in population survival by adaptation score and mutation load. (**D**) Relationship between genetic load and adaptation score of all populations colored by percentage population survival in the 50% reduction treatment. Contour lines represent different levels of population survival percentage. (**E**) Relationship between adaptation score and (**F**) mutation load and percentage survival plotted by treatment along the precipitation gradient. For the same analysis conducted with silique number as fitness metric see **Figure S7,10-11.**

Remarkably, the predictability of fitness differences grew stronger as the treatments became more stressful (**Fig. 2C-E**), and the correlation of scores and survival reached R^2^ = 0.12 in the most extreme 50% rainfall exclusion (*β [SE]* = 4.61 [0.540], *P =* 5.45 × 10^-8^, **Table S6, Fig. S9**). As a positive control of our genomic predictions, we planted trays of accessions based on geographic region alone. We found that our “edge” populations—synthetic populations created directly choosing accessions from North Africa and Southern Europe—show commensurate but slightly lower fitness than genomic-predicted high adaptation score populations (driest treatment: survival _edge pops._= 64% [CI95% = 63% - 73%], **Fig. S8**). While genomic offset and polygenic genomic prediction of maladaptation are often applied to natural populations such as with *Picea glauca* and *Panicum virgatum* ^59,60^, these predictions have not been experimentally tested in accelerated climate change environments ^61–64^. We found these genomic techniques show high predictability of population survival especially in highly stressful and realistic future rainfall reduction.

We then asked whether the genomic adaptation score recapitulates known patterns of adaptive life-history differentiation ^65–67^. Previous studies using this collection of *A. thaliana* accessions performed GWA analyses of fitness from a Spanish outdoor common garden and linked these results to a curated, imputed phenotypic dataset linking fitness with the fast-to-slow growth strategy in *A. thaliana* ^68^. This showed that accessions from more water-stressed regions flowered earlier in spring, escaping impending summer drought, while those from less water-stressed regions grow more slowly and have longer lifecycles^69–71^—mirroring a well-known plant economic spectrum gradient across species ^72–74^ (**Fig 3A**). We associated key phenotypic axes known to be involved in local adaptation ^68^ with the adaptation score from our experiment (**Fig. 3B**), finding a significant correlation with the fast-to-slow principal component axis (PC1, *r = -0.55, P =* 2.2 × 10^-16^), and confirming its relationship with key traits: growth rate, spring flowering time, seed dormancy, root growth, or water use efficiency ^68^. For example, early flowering (*r =* -0.33*, P =* 6.36 × 10^-9^) and high dormancy (*r =* 0.41*, P =* 1.60 × 10^-13^) were more associated with accessions that had high adaptation scores (**Fig. 3A-B, Fig. S13;** importantly these relationships were robust to genomic relatedness and population structure, see **Methods**). By decomposing fitness traits across the 14 precipitation gradient treatments in our experiment, we created principal components of fitness or “fitnotypes” ^75,76^. We found significant co-variation in fitness across treatments in a gradient from wet-adapted accessions to dry-adapted accessions as the major fitness PC1 (PVE 50%) (**Fig. 3D**). These results emphasize that adaptation is a complex, multivariate process shaped by past environments and that genomic prediction approaches can capture a predictive fraction of such past adaptation history.

**Figure 3.**
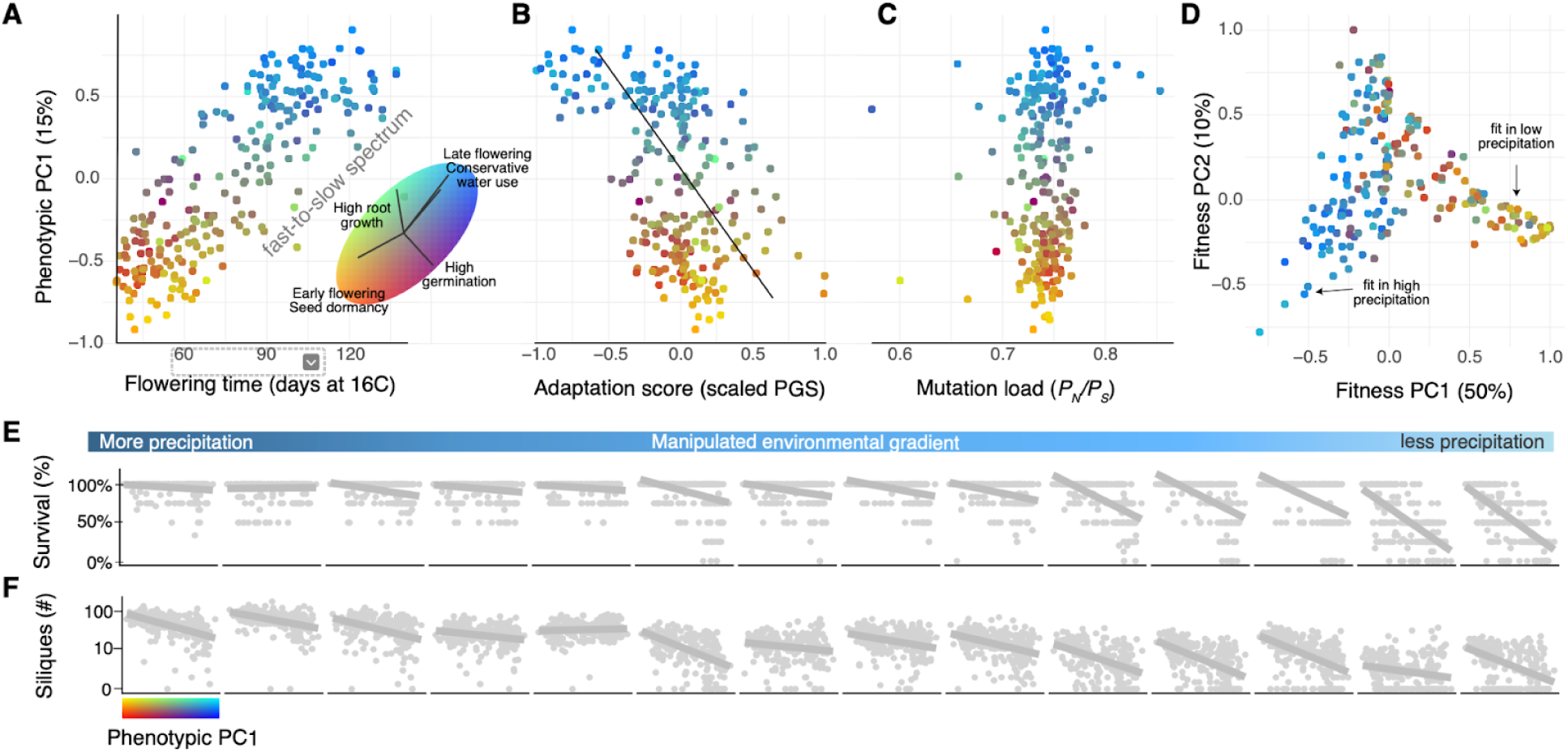
Adaptation score is related to the fast-to-slow ecophysiological adaptation spectrum. (**A**) Axis 1 of phenotypic principal component analysis (PCA) of 12 previously curated traits^68^ compared to flowering time showcases the fast-to-slow adaptation spectrum within *A. thaliana* populations. Colors indicate a bivariate space and ellipsoid shows the main phenotypes loadings with axes scaled to the proportion of variance explained (PC1 15%, PC2 8%). (**B**) Relationship between phenotypic PC1 and adaptation score. (**C**) Relationship between phenotypic PC1 and genetic load. (**D**) Principal component analysis (PCA) of survival data across 14 environmental treatments with phenotypic colors of (A). (**E, F**) relationship between phenotypic axis and survival (E) and silique number (F) across the environmental gradient. The blue gradient bar along the top of D shows the panel order on the precipitation gradient. For the relationship between specific traits and adaptation score and mutation load, see **Figure S13-14**.

### Mutation load synergistically decreases survival probability in extreme stochastic and scarce rainfall environments

To study the role of putatively deleterious mutations in fitness across our climate gradient, we generated several “mutation load scores” by mining whole-genome sequences from the 1001 Genomes. While GWA-based local adaptation scores capture fitness-relevant axes of genetic makeup of populations, natural populations also vary in their composition of rare, potentially deleterious mutations (such rare deleterious mutations are unlikely detectable in GWAs, see discussion in **Text S2**). Using mutation-effect prediction methods, we quantified the ratio of nonsynonymous (e.g. codon missense, stop codon, etc.) and synonymous mutations relative to the genome reference (*P_N_/ P_S_*) (appropriately scaled by overall mutation composition, see **Methods)**. A genome-wide value for *P_N_/ P_S_* of ∼1 reflects inefficient purifying selection purging likely deleterious mutations, which is typical of population histories characterized by bottlenecks and drift, while values of *P_N_/ P_S_* tending towards 0 reflect efficient purifying selection. We found this genome-wide *P_N_/ P_S_* correlates with other clear metrics of likely deleterious variation, including the number of likely loss-of-function gene mutations ^77^, number of singletons, and is robust to recalculation with *A. lyrata* as recent outgroup (**Methods, Figure S5**). Across the *A. thaliana*’s geographic range, it appears that mutation load (*P_N_/ P_S_*) correlates with latitude, likely a legacy of post-glacial expansion that created recurrent bottlenecks as populations migrated north ^78,79^ (*r* =0.32, *P =* 2.2 × 10^-16^) (**Fig. 2A**). Because of this latitudinal trend, we also see mutation load to negatively correlate with annual mean temperature (bio1, *r* = -0.43, *P =* 1.89 × 10^-12^). Experimental populations created with the highest mutation load proxy had significantly lower fitness than experimental populations with low mutation load scores (m_high α_= 43% [CI95% = 36% - 50%, m _low α_ = 60% [CI95% = 51% - 69%, Wilcoxon 2.37 × 10^4^, *P* = 1.78 × 10^-4^, **Fig. 2B**, **Table S4**). As before, the six precipitation treatments with the most extreme rainfall limitation showed the most striking signals, and load scores predicted fitness most strongly in the no-rainfall-addition treatment (*β* [SE] = -7.78 [1.09], *P =* 2.28 × 10^-12^, **Fig. 2C-F, Table S6**), indicating that mutation load impacts fitness primarily toward the drier end of the species’ climatic niche (**Fig. 2F, Table S6**). In addition, unlike adaptation scores, we found a weaker correlation of mutation load score with the trait PC of the fast-slow spectrum (*r = 0.*13, *P =* 2 × 10^-16^, **Fig. 3C, Fig. S14**), in agreement with the notion that deleterious mutations are globally disadvantageous in fitness rather than linked to the same set of traits. Our findings overall show that understanding the relationship between load and future stress environments will have implications for wild populations that may be ill-prepared for a stochastic or intense climate event and may result in fitness declines or extinction ^80; 27,81^.

Theory predicts that both local (mal)adaptation and deleterious load of natural populations should impact their capacity to survive extreme climates, especially as extreme or novel climates may unmask cryptic deleterious variation that was previously not impacting fitness ^36,82^. However, it is still unknown whether deleterious variants cancel out or exacerbate mismatched local adaptation alleles. To better understand how mutation load and adaptation scores interacted to predict fitness in our high stress environments, we tested a logistic regression model to predict survival versus mortality in the most extreme environment of 50% rainfall reduction. We found both mutation load and adaptation score independently had a significant relationship with survival, as well as a synergistic interaction between these two variables on survival (overall GLM *R²* = 0.39, *P_overall_ =* 2.2 × 10^-16^, *P_adaptation score_* = 0.011, *P_genetic load_* = 0.033, *P_interaction_* =0.030)—that is, a high mutation load and low adaptation score synergistically decrease fitness (**Fig. 2E**). Intuitively, this shows that individuals with a *P_N_/P_S_ <* 0.75 and above-median adaptation score *>*0 have >70% chance of survival under an extreme climate-simulated environment (**Fig. 2E**). This significance was not driven by co-variation between scores as they do not appear to be correlated in *A. thaliana* and are accounted for in the model (R^2^ = 0.09, *P* = 0.168, **Fig. 3C**; but see recent contrary example in *Vitis arizonica* ^33^). It is unclear why such an interaction may exist, but possible explanations include epistasis, the multiplicativeness of fitness, or the unaccounted environmental or genomic correlates ^27,83,84^. Regardless of the underlying cause, it is critical to understand that this is not additive—under high stress conditions expected with global climate change, mutational load and maladaptation will synergistically erode population fitness.

Finally we quantified how these detrimental synergistic effects of high mutation load and low adaptation score affect experimental populations in future generations. By allowing the experimental populations to re-sow themselves every spring and re-emerge naturally with fall rainfall, we recorded their population-level survival and fecundity trajectories for a total of 3 years (until 2024) (**Fig. 4A-B**, see **Methods**) in eight of the fourteen treatments that were selected based on their distinct fitness results from the other treatments. The selection pressure from the first year was so extreme that some populations were almost completely lost by the end of year 3 (5/205 trays < 15 individuals by year 3, **Fig. 4**). Unsurprisingly, we found that more stressful precipitation conditions in the first year led to a lower population growth rate, *log*(*λ*), overall across populations (log linear *N* regression: *β*[SE] *=* -0.19[0.003]*, P* < 2 × 10^-16^, *R²* = 0.70). When we grouped the populations on whether they had a genetic makeup of “low-risk” (i.e. either high adaptation score, low mutation load, or both) or “high-risk” (i.e. low adaptation score, high mutation load, or both), we found that low-risk genetic makeup populations had significantly higher growth rate than those with high-risk genetic makeup across all treatments (overall t-test = 12.9, P = 2.2 × 10^-16^) (**Fig. 4C**, see **Methods**). Further, the high-risk genetic makeup populations showed the highest stochasticity of growth rates across replicates in the most water-stressed conditions (*F* test, variance ratio = 12.9, *P <* 2 × 10^-16^) (**Fig. 4C**), which suggests the reduced probability of high-risk populations to be "rescued" if the conditions became more favorable. These results emphasize that conservation goals must target both large and locally adapted populations in order to fully protect and rescue populations under future climates ^85–87^.

**Figure 4.**
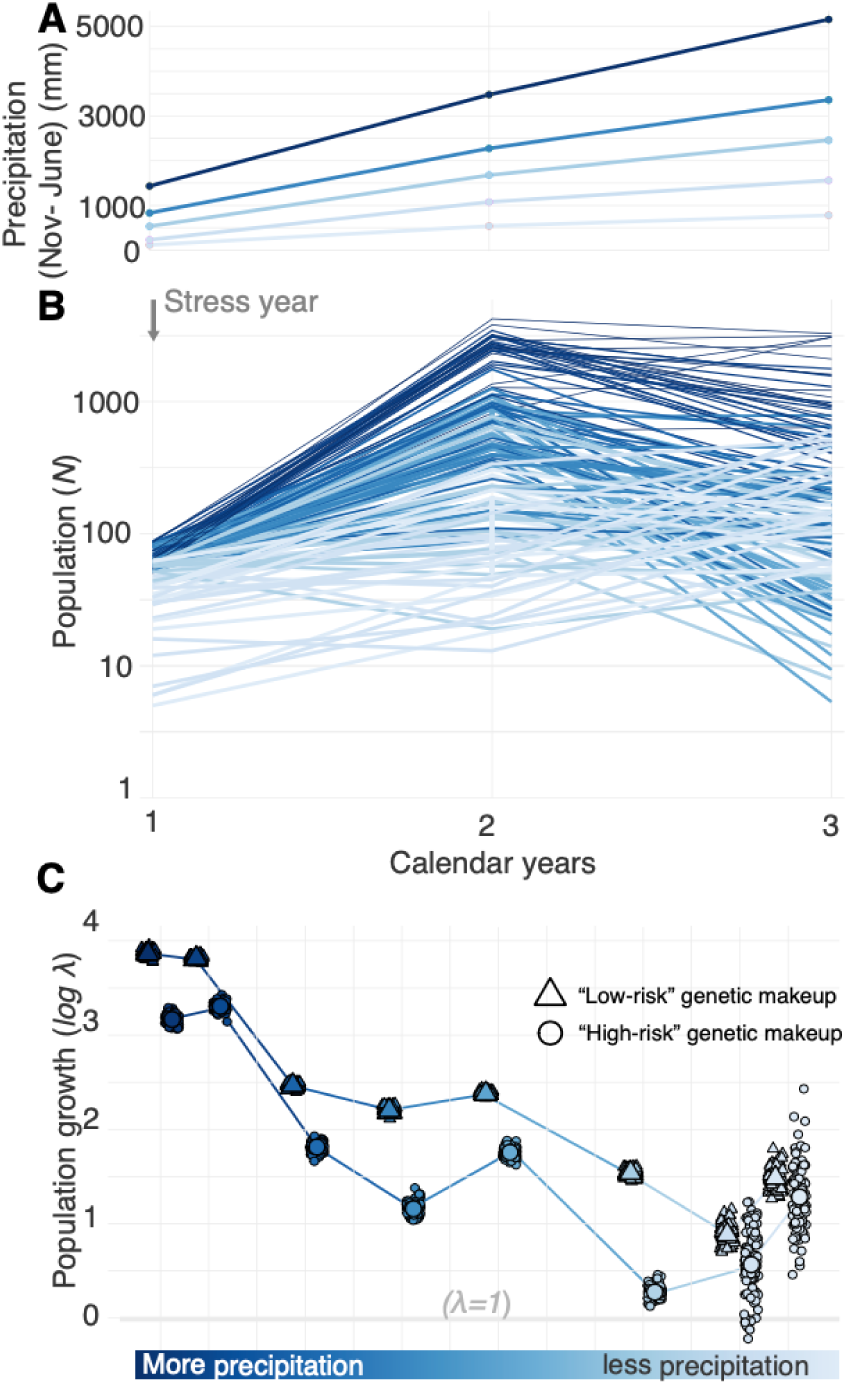
Predicted low-risk genetic makeups have increased rescue trajectory after climate manipulation. (**A**) Precipitation received over the three years of the experiment includes the natural rainfall in addition to the respective rainfall addition or removal for each treatment. Darker blues indicate a higher precipitation addition and light blues indicate the lower precipitation, natural, and exclusion treatments. (**B**) Number of surviving individuals per experimental population across years and precipitation treatments. (**C**) Population growth rate over three years (*logλ*) comparing experimental populations with predicted low-risk genetic makeup (high adaptation score or low mutation load) and high-risk genetic makeups (low adaptation score or high mutation load). Dots represent 100 bootstrap resamples.

## Conclusion

With expected declines in precipitation in the Mediterranean and more stochastic rainfall in European regions ^88^, it is projected that many plant species will require undergoing evolutionary adaptation ^89^. The patterns of locally adaptive genetic composition of *A. thaliana* across the species’ distribution aligns with the hypothesized idea that equatorial edge populations of temperate species contain more thermic and xeric adaptations ^8^. This labeled trailing edge populations as high priority for conservation and disproportionately valued them for the long-term survival of a species. In addition, our results add to the prediction that the fitness effects of mutation load increases with latitude, likely due to post-glacial northward migration and bottlenecks, while it is lower in equatorial edge populations—a pattern now suggested across many plant species ^29,31–33^. Edge populations with synergistic genetic makeups containing locally adaptive alleles and reduced mutation loads may be the best reservoirs for population rescue, while post-glacial populations may be most at-risk. Our research shows that overlapping high-risk genetic makeups are a hidden vulnerability in the conservation of species and will be exacerbated by environmental change.

## Acknowledgements

We are grateful to everyone who volunteered to make this project possible from start to finish. We thank the Moi Lab and Consortium authors for comments on the manuscript. We would like to thank Dr. Tad Fukami, Dr. Molly Schumer, and Dr. Oliver Bossdorf for feedback on the experimental design and analysis. We thank Ismael Villa, Theo Van De Sande, and Johnny Materassi-Shultz for their support and expertise with the field and Carnegie Institution for Science facilities. This work was done in part on the lands of the Muwekma Ohlone Tribe, which was and continues to be of great importance to the Ohlone people, and of the Anishinaabeg – the Three Fires Confederacy of Ojibwe, Odawa, and Potawatomi peoples. Thank you to all the volunteers in and out of the consortium who made this project possible including: Evan Parris, Devaki Bhaya, Flavia Bossi, Adrien Burlacot, Joel Erberich, Weichao Huang, Jacob Irby, Sean Kyota-Waterton, Patti Lang, Rosa Martinez, Maike Morrison, Veronica Pagowski, Andrea Ramirez, Julian Regalado, Bijie Ren, Andrés Reyes, Ruben Shrestha, Ken Thompson, Mackenzie Urquhart-Cronish, Willian Viana, Maria Viteri, Macy Vollbrecht, and Clemens Weiss.

## Additional information

### Author contribution

L.L., M.R.R. and M.E.-A. designed the project. L.L. and M.R.R conducted the field preparations and experimental work with assistance from the field team. L.L. conducted analyses with consultations with M.R.R and M.E.-A. L.L. prepared the manuscript draft. L.L. and M.E.-A. wrote and edited the final manuscript.

### Data and code availability

All code required to reproduce the analyses presented in this study is openly available at https://github.com/lleventhal/ClimateEvo_CommonGarden.

## Methods

### 1. Field experiment design

We created a gradient of precipitation treatments that represent both the frequency and abundances of rainfall experienced throughout the natural range of *A. thaliana.* We designed a common garden in a 7 x 17.4m plot at the Carnegie Institution for Science, in Stanford, California (US) (37.428N, -122.179W) (**Fig. 1A**) with 14 unique combinations of precipitation frequency and abundance as our experimental treatments. We equipped this plot with an aerial irrigation system that ran along nine rows with three sections, resulting in a total of 27 independent irrigation zones (**Fig. S2**). We controlled the irrigation system (Hunter HCC 32 Station Wifi Indoor/Outdoor Plastic Controller (HCC 832-PL) (Hunter Industries)) through bluetooth connection slightly off of the field site. This irrigation controller allowed for water to pump through tubes that were mounted 1m above the trays of plants and were disseminated through six circular sprinklers per section oriented towards the ground to simulate rainfall.

We selected three abundances and four frequencies of watering following a full factorial design across the controller zones. We chose abundances that represented a high (1200mm), average (600mm), and low amount (300mm) of rainfall received in the reproductive season of the *A. thaliana* ecotypes in their native range. We also prioritized varying the frequency of precipitation because the frequency, along with abundance, has been proposed as an important determinant of plant survival ^51^. We selected watering every other day, once a week, every other week, and once a month as our four frequencies. To understand the effects of the natural frequency and abundance of California rainfall, we had one treatment that had no water manipulation whatsoever, just the natural rainfall experienced in our region in that year. Finally, to reduce the rainfall abundance but not alter the frequency of the natural rainfall, we built rain exclusion shelters to reduce 50% of the rainfall by diverting it from the pots (**Fig. 1A, Fig. S2**). All precipitation treatments were replicated twice in this experiment except the treatment of monthly low rainfall because of its similarity to California’s natural weather and space constraints in the field site.

We planted all accession seeds in 30cm x 50cm trays that had 60 6cm x 6cm pots each (**Fig. 1C-D**). We filled the pots with Nutrient Ag Solutions PROMIX PGX Biofungicide Plug & Germination mix. In total, we prepared 243 trays for our accessions analyzed in this paper. To prevent seed carryover after sowing, we gently watered the trays for two weeks until germinants were established. After germination was established, we thinned each pot to have a single plant. We considered each tray with ∼60 plants each to be a population, and population means were measured by taking averages across the contained individuals.

### 2. Accession selection

We planted a diverse panel of accessions in curated populations. For these experimental populations, we selected 245 accessions of *Arabidopsis thaliana* from the 1001 Genomes Consortium to maximize geographic spread, lower genetic redundancy, and avoid ecotypes with lower genome data quality, and for which we already have previous fitness data in two native range common gardens ^18^. In our study, an experimental population is a subset of selected accessions that have our “target” genomic characteristic in common including mutation load and polygenic score for local adaptation (see below).

The 245 accessions had a range of genetic makeups, and were selected based on criteria such as mutation load, adaptation score, and proximity to Northern Africa (see methods sections 2.1 and 2.2), and we replicated each accession at least four times per treatment, culminating in 14,240 individuals). In total for this study, we created five experimental populations of different genetic makeups, and each synthetic population was housed in its own tray. We planted accessions in synthetic population groups in a tray in a randomized block design rather than a fully randomized design because the synthetic population represents in this experiment a similar genetic makeup.

The accessions included in each synthetic population (high and low mutation load, high and low adaptation score, and geographic range edge) were not mutually exclusive (e.g. an accession with a high adaptation score for drought survival may also be on the southern edge of the range). In the case of an accession overlapping categories, we included it in both populations as the intention of this experiment was to choose genetically motivated populations. Each population was replicated twice within each treatment block, and as there were two blocks per treatment (except monthly minimum). We planted all of the accessions in a randomized complete block design. We randomized the location of the accessions in each tray and each tray within its replicate block. Finally, we randomized the treatment blocks within the common garden.

#### 2.1 Quantifying mutation load

We calculated mutation load by counting the number of nonsynonymous polymorphisms per nonsynonymous site (*P*_N_) divided by the number of synonymous polymorphisms per synonymous site (*P*_S_), resulting in the ratio of of nonsynonymous to synonymous mutations within the genome (*P*_N_/*P*_S_). We found *P*_N_ and *P*_S_ relative to the *A. thaliana* reference genome, Col-0, using snpEff for annotation ^28^ and bcftools for filtering after annotation ^90^. We counted annotations as *P*_N_ if it led to a change in a protein coding region (see **Table S7**), while we counted an annotation as *P*_S_ if it did not create a change in a protein coding region. We selected 60 ecotypes that had the highest *P*_N_/*P*_S_ for the high mutation load populations and 54 ecotypes that had the lowest *P*_N_/*P*_S_ for the low mutation load populations. Although the goal was 60 accessions per population, sometimes seed limitations made the population numbers smaller.

To verify the validity of our mutation load proxy (*P*_N_/*P*_S_), we used a congeneric reference, *Arabidopsis lyrata* v.2, to minimize potential reference bias due to genetic relatedness and population structure. We first aligned the *A. thaliana* Col-0 reference genome to the *A. lyrata* v.2 genome using Anchorwave with default parameters ^91^, generating a high-confidence cross-species coordinate map. We transferred 1001 Genomes variants calls to the *A. lyrata* genome coordinates and filtered variants for quality using criteria equivalent to our original pipeline (e.g. minimum depth, mapping quality threshold). We then recalculated *P*_N_/*P*_S_ for each accession using the same functional annotation strategy with snpEff as with Col-0. This allowed us to compare the *P*_N_/*P*_S_ distributions relative to a more genetically distant reference, provisioning a robust test of our mutation load estimate and evaluating the sensitivity of our approach to reference genome choice (**Fig. S5**).

#### 2.2 Calculating polygenic score for local adaptation

To quantify a polygenic score for local adaptation to drought conditions, we utilized polygenic scores (PGS) and counted variants associated with survival in drought conditions in a previous drought experiment including 515 accessions ^18^ by measuring effect sizes estimated from genome-wide association studies (GWAs; ^92,93^. We used a linear mixed model using GEMMA 0.98.3 ^94^ to estimate the effect sizes of each SNP (sequences from the ^46^) to identify which SNPs most associate with survival in this drought treatment. We calculated PGS using the effect sizes estimated for each SNP, the significance of each SNP, as well as covariate data for the selected samples, including relatedness. We calculated PGS following ^47^ while accounting for linkage disequilibrium and population structure using the software PLINK v2.00a2.3 ^95^. We tested the polygenic scores with different sets of SNPs that varied in significance cut off from 10^-3^ to 10^-9^, and we selected the association threshold of 0.001. For the high and low adaptation score populations, we selected 57 accessions with highest PGS and 59 accessions with the lowest PGS (hereafter the high and low adaptation score populations).

### 3. Sowing protocol and quality control

On November 16, 2021, we synchronously sowed 14,232 *A. thaliana* pots with 245 accession. All volunteers followed the same protocol of seed sowing to avoid contamination. Prior to planting, we individually aliquoted seeds into 2ml Eppendorf tubes and labeled them with the accession identification. We randomly placed the eppendorf tubes into cardboard freezer storage boxes with compartments, matching their field placement orientation in the 6 x 10 grid trays. We recorded the position of each accession within each box. Volunteers received a prepared box for a random location in the field and sowed the seeds by carefulling dispensing them from the tubes, maintaining a height of 1-2cm above the soil surface to avoid contamination. We recorded any errors in planting or potential contamination, which was used to eliminate those replicates in downstream analyses.

### 4. Data collection

#### 4.1 Fitness and phenotypes

*Germination.* After two weeks of watering after sowing, we surveyed all of the pots for germination. If they did not germinate, we recorded the position and marked the pot with a tag.

*Flowering time.* We surveyed the plants for flowering time every one to three days depending on the timing of the flowering season from January to June 2022. We considered the first day of flowering in *A. thaliana* as the first day petals were visible emerging from the sepals. Upon flowering, we put a blue tag in the pot to eliminate redundancy in counting and improve the surveyor’s efficiency. We calculated flowering time as days since sowing to the first day of flowering.

*Survival.* We recorded survival three to four times a week. If an individual was dried and shriveled, it was counted as dead. Any individuals that died before flowering were counted as dead. If an individual flowered, and the flowers never produced siliques and the rosette and inflorescence were dry and shriveled, we recorded it as dead. We recorded survival for all 14,232 plants.

*Fecundity.* We recorded the silique number for 12,281 plants by counting the number of siliques in the field using a hand counter, although 2,604 of these plants survived to flowering but did not create any siliques, and we recorded a silique number of zero. The remaining 1,951 plants did not have a silique number but were recorded as NA because they died before flowering. The number of siliques that a surviving *A. thaliana* plant produces is a suitable proxy for the number of seeds, as the average *A. thaliana* silique has about 20 seeds ^96^.

#### 4.2 Meta data sensors

We measured temperature (°C), soil moisture (%), humidity, soil fertility (µS/cm), and light intensity (lux) using Huacaocao Plant Monitors (Beijing Huahuacao Technology Co., Beijing, China) and temperature and air humidity using iButtons (iButton Link, WI, US). We had at least two soil moisture sensors per precipitation zone and one iButton per zone. Through these sensors, we were able to monitor the treatments, evaluate heterogeneity across the garden, and compare variation within and among precipitation treatments.

### 5. Statistical analyses and methods

All statistical analyses were conducted using R version 2024.09.1.

#### 5.1 Testing effect of frequency and abundance of precipitation on fitness

To test the relative effects of rainfall frequency and abundance on our two fitness measures, survival and fecundity, we fit a Bayesian generalized linear mixed model using the MCMCglmm package in R (ref. ^97^. For the survival model, our response variable was the binary outcome of survival (0 or 1), thus we used family = categorical, meaning a categorical distribution with a logit link. The model was structured to have an intercept-only fixed effect and had our focal random effects of rainfall treatments (frequency, abundance, frequency x abundance) as well as random effects to account for experimental design including tray position, treatment location in field, and coordinates in the tray. The silique response variable model used family = poisson with log link, and had the intercept-only fixed effects and the same random effects, with an additional term of observation-level random effect to account for overdispersion of the poisson distribution. The priors were weakly informative (fixed effects: diffuse normal [μ=0, σ²=1×10⁸], random effects: Inverse-Wishart (V=1, ν=0.002), poisson residual variance fixed at 1)), and the model ran for 13,000 iterations with 3,000 burn-in (23% discarded), 10-iteration thinning (retained 1,000 samples), and latent variables (pl=TRUE) and predictions (pr = TRUE) stored. We checked effective sampling size was > 1,000 for all parameters, and autocorrelation was < 0.1 at lag 10. Additionally, we did a visual inspection of trace plots for mixing.

After fitting the models, we extracted the posterior distributions of the variance components associated with each random effect (frequency, abundance, and their interaction). For each effect, we summarized the posterior distribution by reporting the posterior mode as the effect size, along with the 95% credible interval (the 2.5th and 97.5th percentiles of the posterior). This approach quantifies both the magnitude and direction of each effect on the fitness measure, accounting for uncertainty.

Further, we tested the relationship between home environment precipitation stochasticity and performance across our varying stochasticity environments. To measure home environment stochasticity by month, we downloaded the TerraClimate dataset ^98^ from 1958 to 2017 and extracted precipitation at all accessions’ location of origin. To quantify between-year variability we quantified per month Coefficient of Variation of amount of precipitation across the entire time series.

#### 5.2 Testing the role of population background on fitness across treatments

We used a Wilcoxon rank sum test to determine if paired populations (high vs. low mutation load, high vs. low adaptation score) had significantly different fitness in drought conditions. After comparing experimentally paired populations, we studied the relationship of all accessions’ deleterious load and adaptation score with fitness using a Pearson correlation and a linear mixed model. In this linear model, we filtered the data by experimental treatment, and then used the genomic measure of interest (e.g. adaptation score, deleterious load) as predictor variable and fitness as response variable. We also conducted a Pearson correlation to find the relationship between adaptation score and deleterious load. To determine the impact of adaptation score and load separately and their interaction on fitness, we used a linear model where the response variable was proportion survival (filtered by treatment) and ranked adaptation score and inverse load (i.e. 1-load, in order for the hypothesis that a lower load is positively related with fitness) were fitted as fixed effects with an interaction component. For this analysis, we did a split normalization of adaptation score around the median, which normalizes values above and below the median separately.

#### 5.3 Calculating the heritability of fitness

We used a hierarchical Bayesian generalized linear mixed model using the MCMCglmm package in R to estimate broad-sense heritability (*H^2^*) of survival probability and fecundity across all treatments combined. We created an intercept-only model, meaning we included no fixed covariates, but we included random effects of accession *id*, tray, zone within the field, and position within the tray. The *id* effect is informed by the kinship matrix created in GEMMA 0.98.3 ^94^ (using command "-gk”)as variance-covariance of the random effects following standard MCMCglmm approaches. The survival model was fit using a binomial family (i.e. Binomial in MCMCglmm nomenclature) because the survival data is binary while the fecundity model was fit using a poisson family as it is count data and right skew distributed. We had default weakly informative priors and had 13,000 iterations (burn-in = 3,000, thin = 10), yielding 1,000 posterior samples. We assessed the model for convergence with trace plots and confirmed an effective sample size (ESS > 1,000) for all parameters. The output of this model was the posterior mean of *H^2^* with 95% highest posterior density intervals. This model robustly quantified genetic contributions to our traits (survival and fecundity) while accounting for experimental design structure.

### 6. Census of populations for three years

In the second and third year of the experiment, we reduced the experiment size from 14 treatments occupying 27 zones to eight treatments occupying 16 zones and 128 experimental populations. This reduction was motivated by the redundancy in the fitness measurements from the first year and due to time and effort constraints. The census of individuals and siliques per experimental population (i.e. tray) in the 2nd and 3rd year of the experiment was conducted based on a sampling approach and inference of the total population-level value as follows: After flowering commenced, we counted how many pots had at least one plant, and then from this subset randomly selected three pots in the sixty pot trays using a random number table. Within these three pots we counted all the individuals and their number of siliques for each individual. With this we obtained a robust average of individuals per pot, and siliques per individual, which we extrapolated to the population level multiplying by all pots with alive plants and all plants with fruits. To assess the robustness of this approach, we cross-validated these predictions counting every individual and silique of 5 random trays across the environmental gradient, which indicated the true population values were always within the 80% confidence interval of the estimator.

Note the timing of the second year’s growth season (fall 2022 – spring 2023) coincided with an atmospheric river of rain in northern California, and the third year (2023–2024) also was characterized by high and frequent rainfall, thus the first year is considered the stress year (**Figure S1**). Years two and three therefore monitor the effects of a single stress year plus treatment interacting with the genetic background of populations over time.

## Supplemental Appendix

### Supplemental Text

#### 1. Thoughts on local adaptation score and links to population structure

The local adaptation score was conducted following standard protocols in the field using effect size estimated in a GWA of survival in 515 accessions and combining effects accounting for population structure and LD in PLINK to project to a subset of 245 accessions (see Methods). If we had perfect knowledge of causal alleles (whether SNPs or structural variants) in a highly heritable trait it would be straightforward to make predictions of a phenotype. Even though we show the polygenic score method has predictability, we do not claim it has perfect knowledge of adaptive loci, nor is this required to achieve predictability ^99^. Most likely our GWA method is identifying linked SNPs to adaptive haplotypes that are then used for prediction. In addition, because a trait like survival under drought in common gardens is aiming to capture local adaptation to such an environment, it is likely that survival is strongly linked to population structure because local adaptation and population structure are intricately related ^18,100^. Precisely population structure is predictive of local adaptation, and researchers may leverage this for short-term predictions. For example, in application for conservation, Lotterhos *et. al* ^101^ found in complex spatially-explicit local adaptation simulations that predictability of fitness and mal-adaptation was predicted using genome-wide composite scores better even than with only top adaptive alleles ^102^. Here and other experiments ^18,58^, we show global panels of populations of a species capture general species-wide patterns that are predictive. However, while likely GWA trained in a global panel of 515 or 240 accessions may be more readily able to predict at smaller scales (i.e. variation within a regional panel) than the other way around, it is still unclear whether small differences in fitness of multiple highly related individuals will be well-predicted. It will remain to be tested in future iterations of our common gardens.

With this reflection, we ensure that we correct for population structure when we conduct correlations among different genomic metrics, such as adaptation score and mutation load, climate, or phenotypes. To do this, we fit major axes of population structure using 3 genomic PCs as fixed effects in regressions or through incorporating a kinship matrix as a random effect model, when appropriate.

#### 2. Various mutation load approaches and population structure

Although *A. thaliana* is a highly self-fertilizing plant (with variable 3–16% outcrossing rates ^17,103,104^), and it is often assumed natural populations efficiently purge deleterious recessive variants ^105,106^, whole-genome sequencing has revealed large variation in past bottlenecks of natural populations ^46,107^, with various levels of low frequency, likely deleterious mutations (stop codon, missense, etc.) ^30^. Such deleterious mutations are unlikely to be detectable in GWA if they are at very low frequency ^53^), thus there are several methods to measure mutational load. Including the number of loss-of-function genes, number of singletons, etc. Here we focus on polymorphisms that change an amino acid in the protein coding sequence in the genome.

A common method when quantifying the efficiency of purifying selection (and thus purging) is the nonsynonymous-to-synonymous substitution rate in genes across species, *d_N_/d_S_* ^15,108^. A *d_N_/d_S_* > 1 indicates positive selection in likely adaptive mutations, while *d_N_/d_S_* < 1 indicates purifying selection predominates eliminating nonsynonymous variants likely deleterious. The combination of substitutions between species with information about polymorphism *P_N_/P_S_* within a focal species is the basis of the McDonald-Kreitman test ^109^. When studying only within-species variation that may indicate differences in populations in drift or purifying selection, we can use *P_N_/P_S_*, the proportion of nonsynonymous-to-synonymous polymorphisms genome-wide in an individual part of a rangewide set of individuals. *P_N_/P_S_* is appropriately scaled by SNP discovery (total synonymous or nonsynonymous SNP sites across all accessions in the analysis) to reduce bias by accounting for variation in SNP detection rates across accessions. *P_N_/P_S_*can then be used as a genome-wide indicator of the efficiency of purifying selection effect of selfing *P_N_/P_S_* ^29^). At the genome-wide level, we may then expect that pN/pS values are close or below one, where one would indicate a population accumulates mutations mostly due to genetic drift, whereas values below one, suggesting different strengths of purifying selection acting in natural populations.

Mutation load is likely linked to population structure because mutation load is accumulated through population demographic processes that cause genome divergence across populations. By creating these experimental populations are “artificial” or “synthetic” populations, as no population exists in the wild with this high level of likely mutation load but diverse genotypes. We could in principle try to identify a single highly bottlenecked population of *A. thaliana* and use multiple individuals within that population for our experimental populations, but this was not the goal of our synthetic population, which aim to be representative of different genotypes across the species range with only the high mutation load variable in common. As before, because we recognize population structure will be linked to the accumulation of deleterious variants (**Fig. 2**), we account for population structure when correlating mutation load with traits or others by controlling by latitude, using 3 main genomic PCs, or the kinship matrix of these accessions random effect models when appropriate for the model and question.

## Supplemental Tables

**Table S1.** Summary of π diversity by population

| **Population** | **Genome $\pi$/SNP diversity** | **Number of accessions mixed in experimental population** |
| --- | --- | --- |
| low polygenic score | 0.132144 | 59 |
| high polygenic score | 0.143000 | 58 |
| high mutation load | 0.129843 | 60 |
| low mutation load | 0.152778 | 60 |
| edge | 0.137599 | 60 |

**Table S2.**
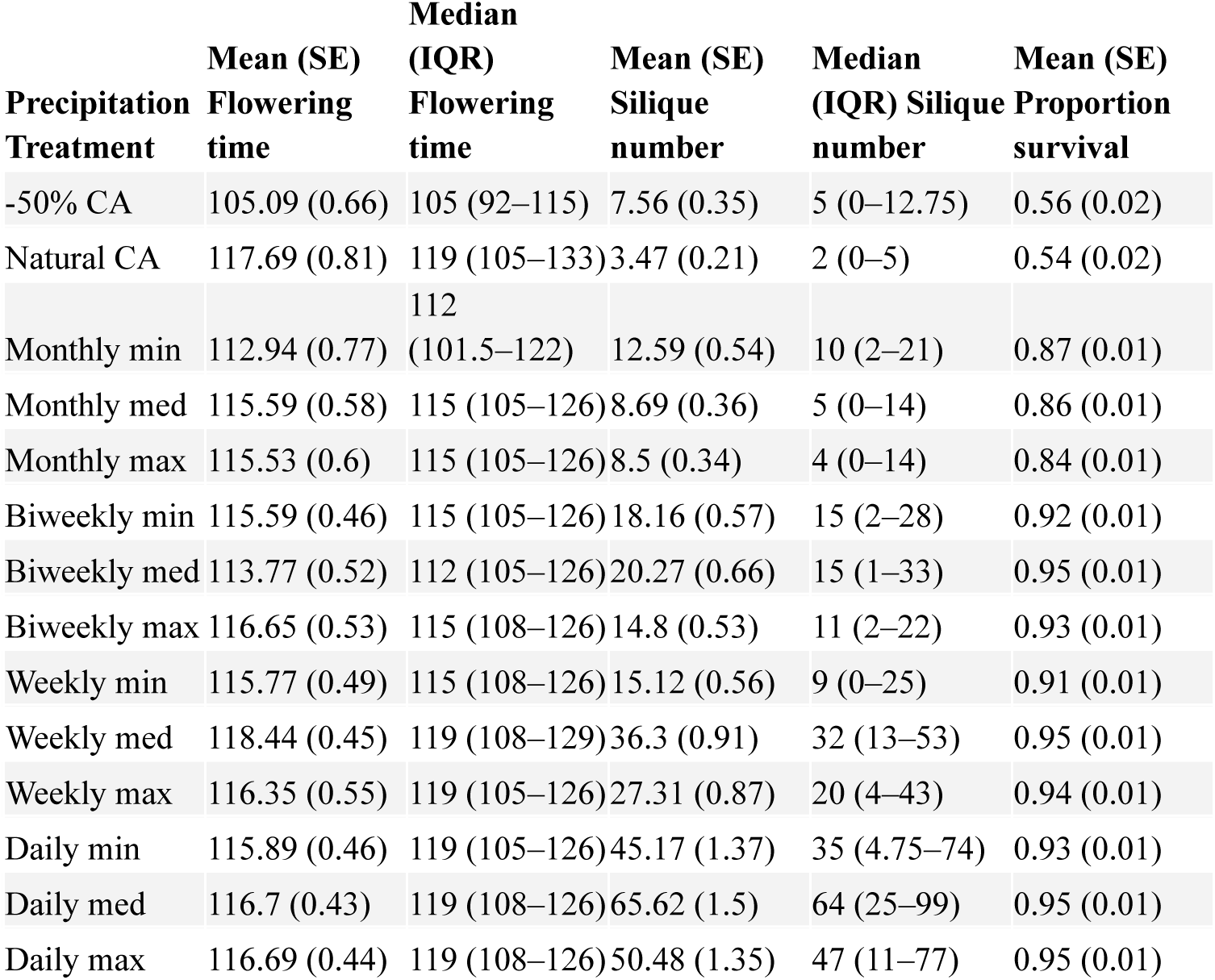
Summary statistics of Flowering time, silique number and proportion survival across treatments.

**Table S3.** Frequency and abundance model and interaction result, percent variance explained (PVE)

| Effect | PVE_Mean | Lower_95CI | Upper_95CI | Fitness |
| --- | --- | --- | --- | --- |
| frequency | 0.461693792 | 0.123540029 | 0.882147289 | survival |
| abundance | 0.031822992 | 1.48E-05 | 0.183062032 | survival |
| frequency:abundance | 0.005460904 | 1.49E-05 | 0.022317181 | survival |
| frequency | 0.36294076 | 0.07519293 | 0.77049541 | silique number |
| abundance | 0.02059689 | 1.66E-05 | 0.08996 | silique number |
| frequency:abundance | 0.01251985 | 9.82E-06 | 0.04630848 | silique number |

**Table S4.**
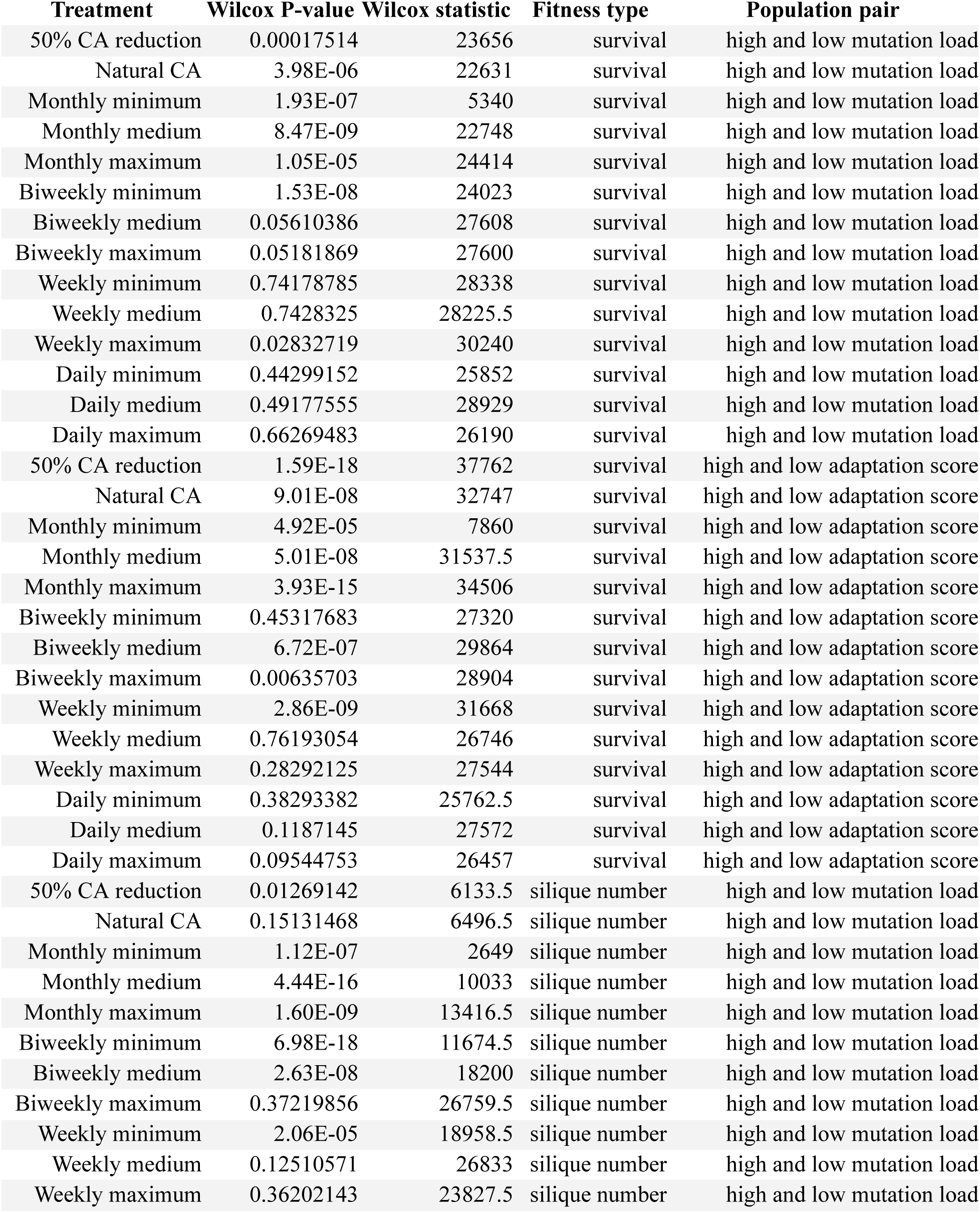
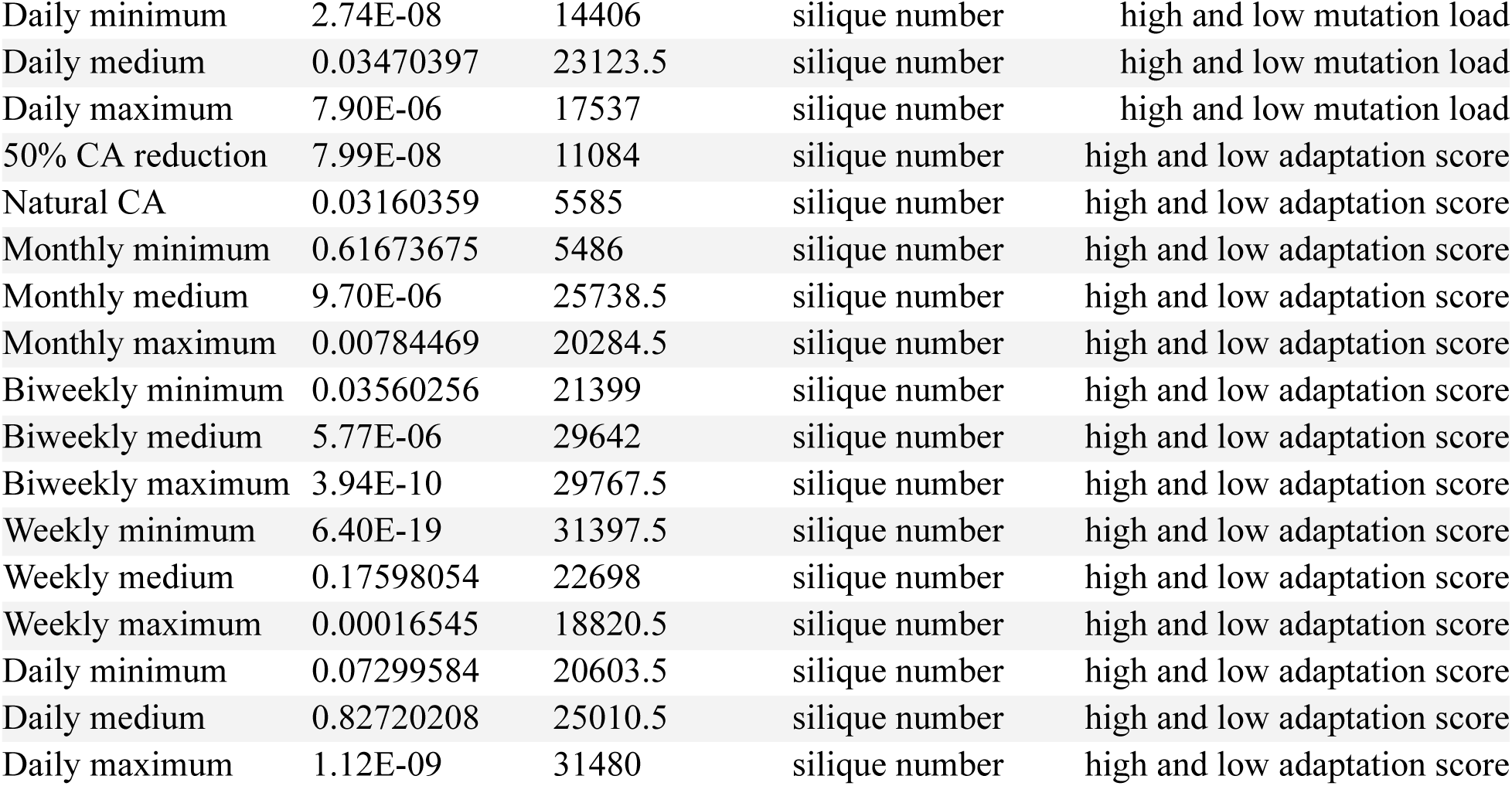
Paired Wilcoxon signed rank z-scores for high and low adaptation score and high and low mutation load populations across precipitation treatments.

**Table S5.** Linear regression output of adaptation score and survival by treatment

| Treatment | slope | SE | P-value | R<sup>2</sup> |
| --- | --- | --- | --- | --- |
| Daily maximum | 1.083941157 | 0.2179547376 | 8.66E-07 | 0.04049741652 |
| Daily medium | -0.3905856693 | 0.2291328433 | 0.08879255325 | 0.004917447215 |
| Daily minimum | 1.72183744 | 0.2844542096 | 2.53E-09 | 0.05875236588 |
| Weekly maximum | 1.02820583 | 0.2648413519 | 0.0001151775339 | 0.02499303088 |
| Weekly medium | 0.2052006661 | 0.2211772926 | 0.3539098789 | 0.00146171886 |
| Weekly minimum | 3.924650384 | 0.4959404278 | 1.26E-14 | 0.09654942826 |
| Biweekly maximum | 2.766103247 | 0.4168681261 | 7.37E-11 | 0.06988417016 |
| Biweekly medium | 3.41132508 | 0.3373304635 | 2.88E-22 | 0.1485861072 |
| Biweekly minimum | 2.64160248 | 0.4121015377 | 2.99E-10 | 0.06541919743 |
| Monthly maximum | 8.13952303 | 0.4700432736 | 1.47E-54 | 0.3384978281 |
| Monthly medium | 7.635191349 | 0.5390466608 | 2.18E-39 | 0.2547223826 |
| Monthly minimum | 5.846779919 | 0.6270543696 | 2.25E-19 | 0.1290034771 |
| Natural CA | 7.223298418 | 0.5580718513 | 6.90E-34 | 0.2220315788 |
| 50% CA reduction | 7.923140727 | 0.568478163 | 2.39E-38 | 0.2486426406 |

**Table S6.** Linear regression output of mutation load and survival by treatment

| Treatment | slope | SE | P-value | R<sup>2</sup> |
| --- | --- | --- | --- | --- |
| Daily maximum | -0.500386868 | 0.3123305 | 0.109458207 | 0.002644585 |
| Daily medium | 0.510007919 | 0.327096226 | 0.119256015 | 0.00235018 |
| Daily minimum | -1.080457925 | 0.370393484 | 0.003616035 | 0.008804061 |
| Weekly maximum | 0.543493688 | 0.388553603 | 0.162202953 | 0.002004734 |
| Weekly medium | 0.13739693 | 0.298743327 | 0.645674497 | 0.00020774 |
| Weekly minimum | -1.249925734 | 0.503966351 | 0.013314253 | 0.006810905 |
| Biweekly maximum | -1.283190697 | 0.415156938 | 0.002054578 | 0.010060937 |
| Biweekly medium | -0.656288171 | 0.369993767 | 0.076421243 | 0.003318371 |
| Biweekly minimum | -2.398864185 | 0.439069507 | 5.96E-08 | 0.03034002 |
| Monthly maximum | -3.469538101 | 0.647400636 | 1.08E-07 | 0.032947286 |
| Monthly medium | -3.166096309 | 0.651265918 | 1.39E-06 | 0.027589027 |
| Monthly minimum | -5.496111729 | 1.002197054 | 6.60E-08 | 0.056523891 |
| Natural CA | -7.779455825 | 1.087022398 | 2.28E-12 | 0.074421002 |
| 50% CA reduction | -5.760246974 | 1.048354149 | 5.45E-08 | 0.040567871 |

**Table S7.** Nonsynonymous and synonymous mutation annotations.

| Nonsynonymous annotations | Synonymous annotations |
| --- | --- |
| missense variant | synonymous variant |
| splice_region_variant | stop_retained |
| splice_donor_variant |  |
| splice acceptor variant |  |
| stop_gained |  |
| stop_lost |  |
| start lost |  |
| frameshift_variant |  |
| initiator_codon_variant |  |

## Supplemental Datasets

Dataset 1. Accession_information

This dataset includes the unique ids for the 245 accessions used in this experiment, their location, and the local adaptation score and mutation load score used to categorize the accessions.

Dataset 2. Experimental_and_fitness_information

This dataset includes all elements of experimental design and the fitness and phenotypes collected during this experiment. The columns are defined as such:

*Dataset column definitions:*

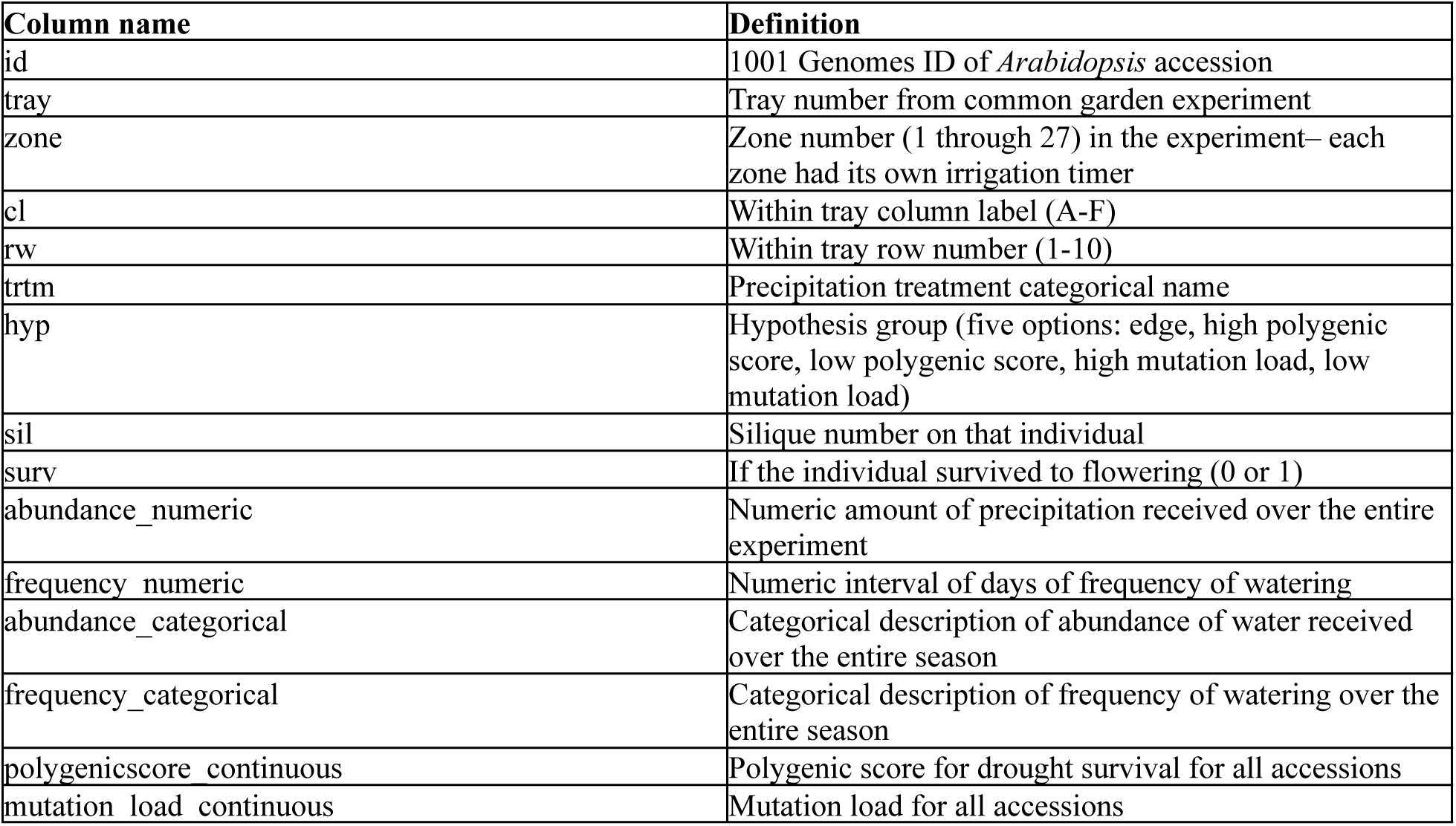

## Supplemental Figures

**Figure S1.**
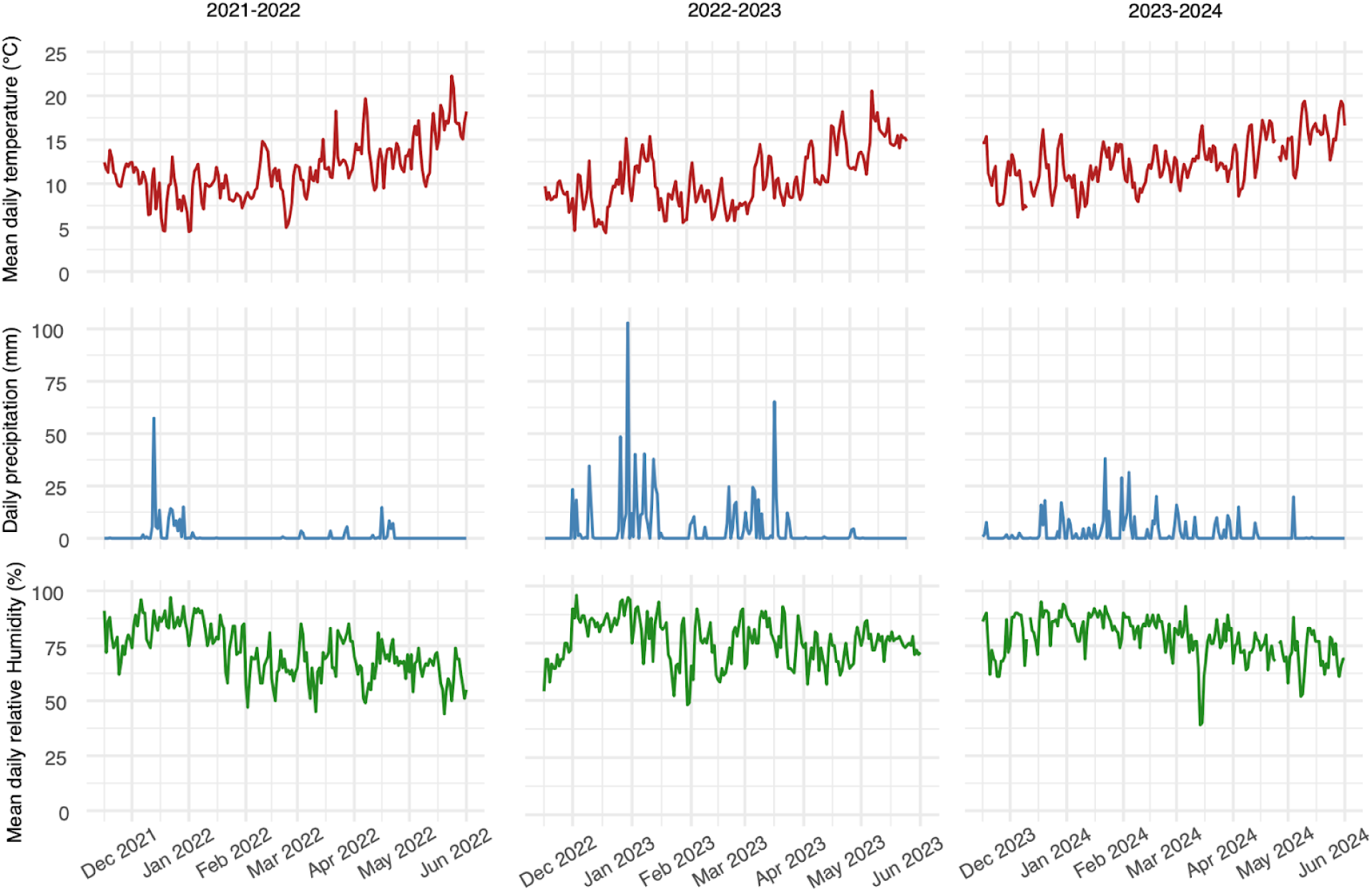
Mean daily temperature, total daily precipitation, and mean daily humidity during 2021-2022, 2022-2023, 2023-2024 field season (mid-November through early June). Data are from Stanford weather station (https://stanford.westernweathergroup.com/).

**Fig. S2.**
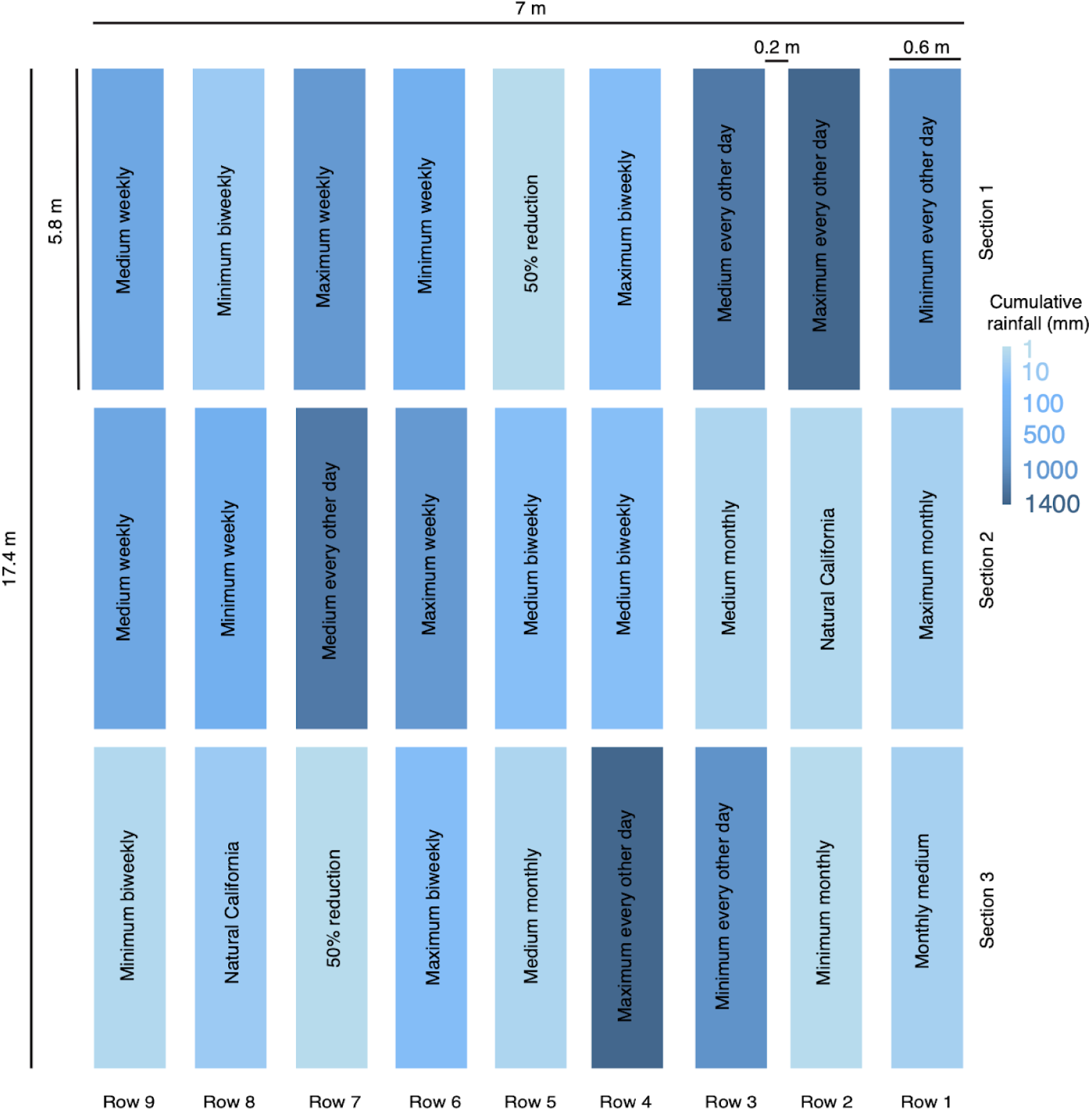
Overview of the common garden site with treatments. The field site was outfitted with 27 independent irrigation timers. We had 14 treatments within this site replicated across the zones. Colors represent the color gradient of drought treatment.

**Figure S3.**
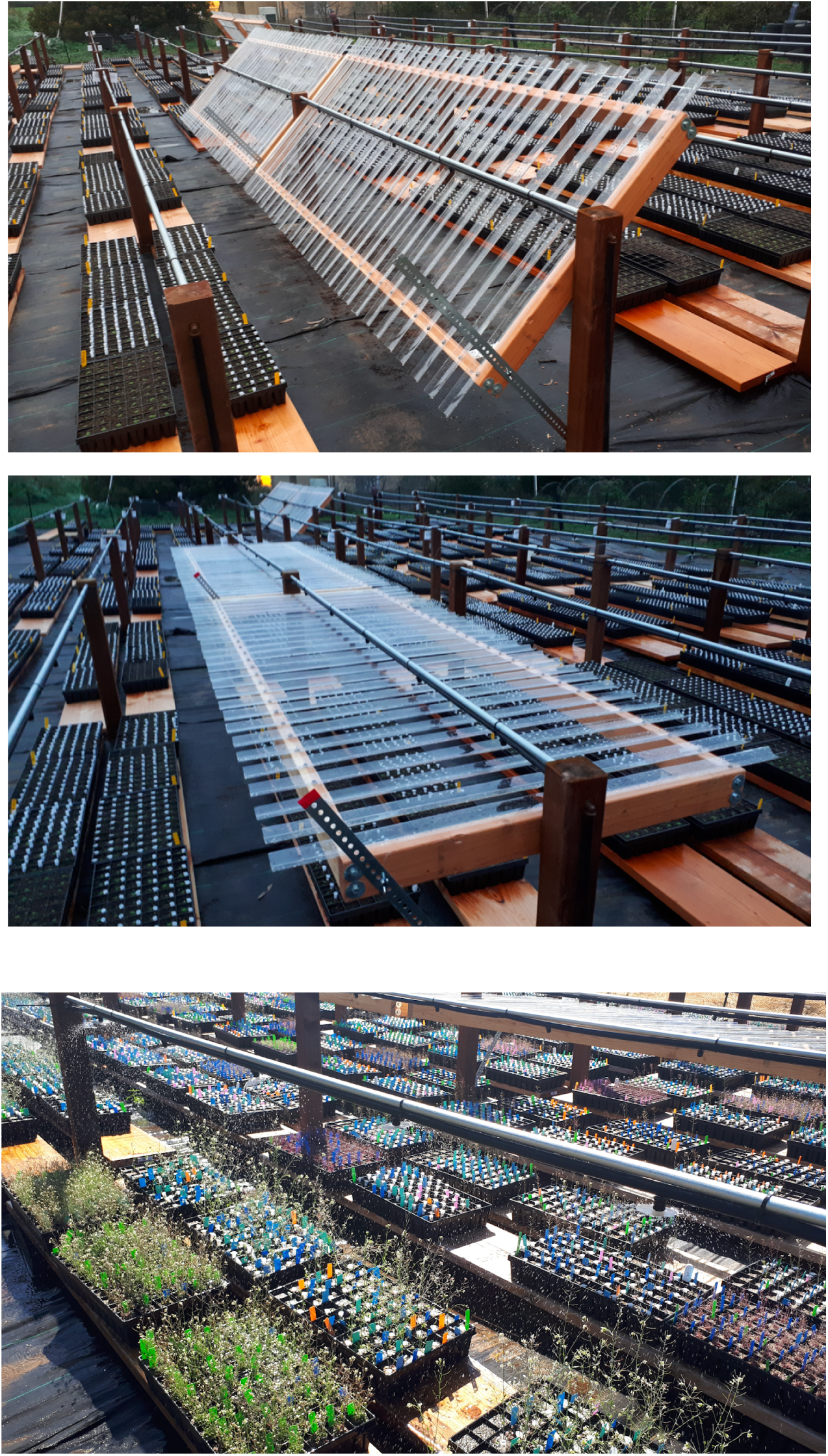
Close up of rainout shelters and aerial sprinklers. Two orientations of rainout shelters, highlighting that they can be rotated in order to reduce rainfall coming from more of an angled direction (top and middle). Photo of aerial sprinklers simulating rainfall in the field (bottom). Photo credit: Lucas Czech.

**Figure S4.**
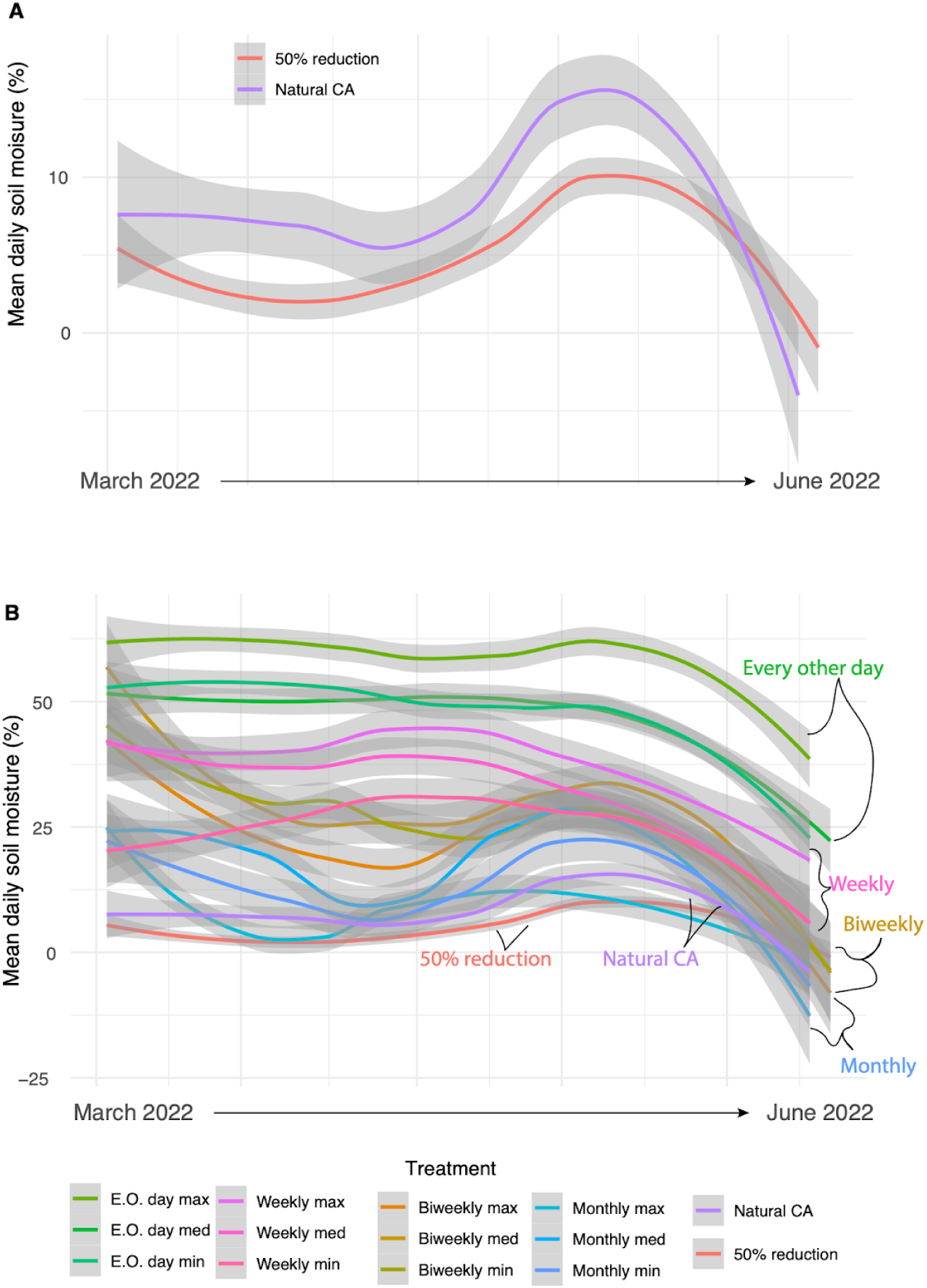
(A) Mean soil moisture across all sensors in the 50% reduction and natural treatments for March through May. The purple line represents the no-additional rainfall treatment, while the red line represents the 50% reduction treatment. Until the pots were completely dry, in about May, the 50% reduction treatment recorded approximately 50% of the soil moisture as the Natural CA treatment. (B) Soil moisture data across all fourteen treatments. Frequencies are grouped by color where greens are every other day, pinks are weekly, oranges are biweekly, blues are monthly, and purple is natural CA no addition, and red is 50% reduction.

**Figure S5.**
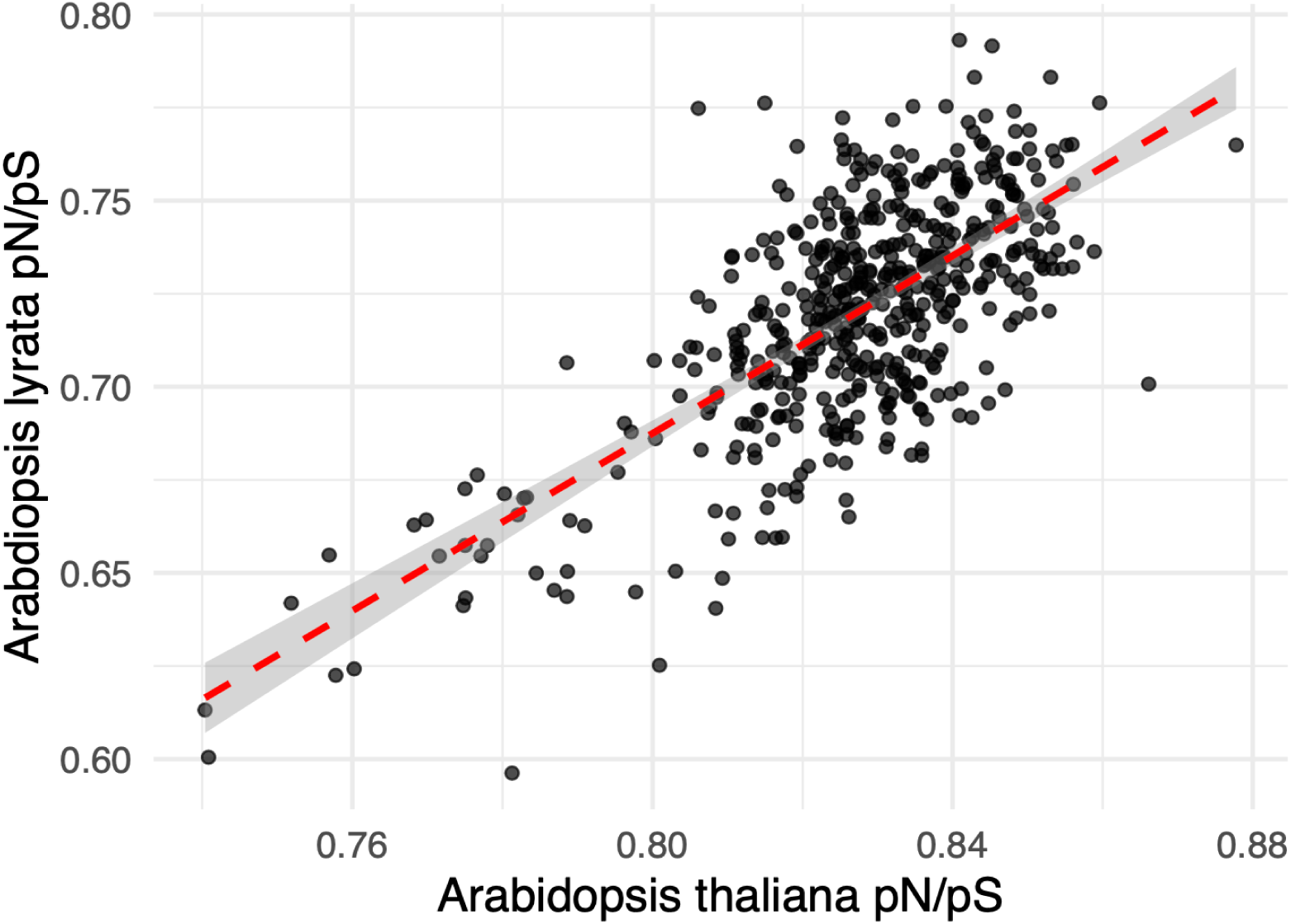
*pN/pS* for Arabidopsis thaliana using Col-0 as the reference for SNP annotations (x-axis) and an outgroup, *Arabidopsis lyrata*, as reference for SNP annotations (y-axis) (R = 0.70, 95%CI = 0.65-0.74, p = 2.2 × 10^-16^).

**Figure S6.**
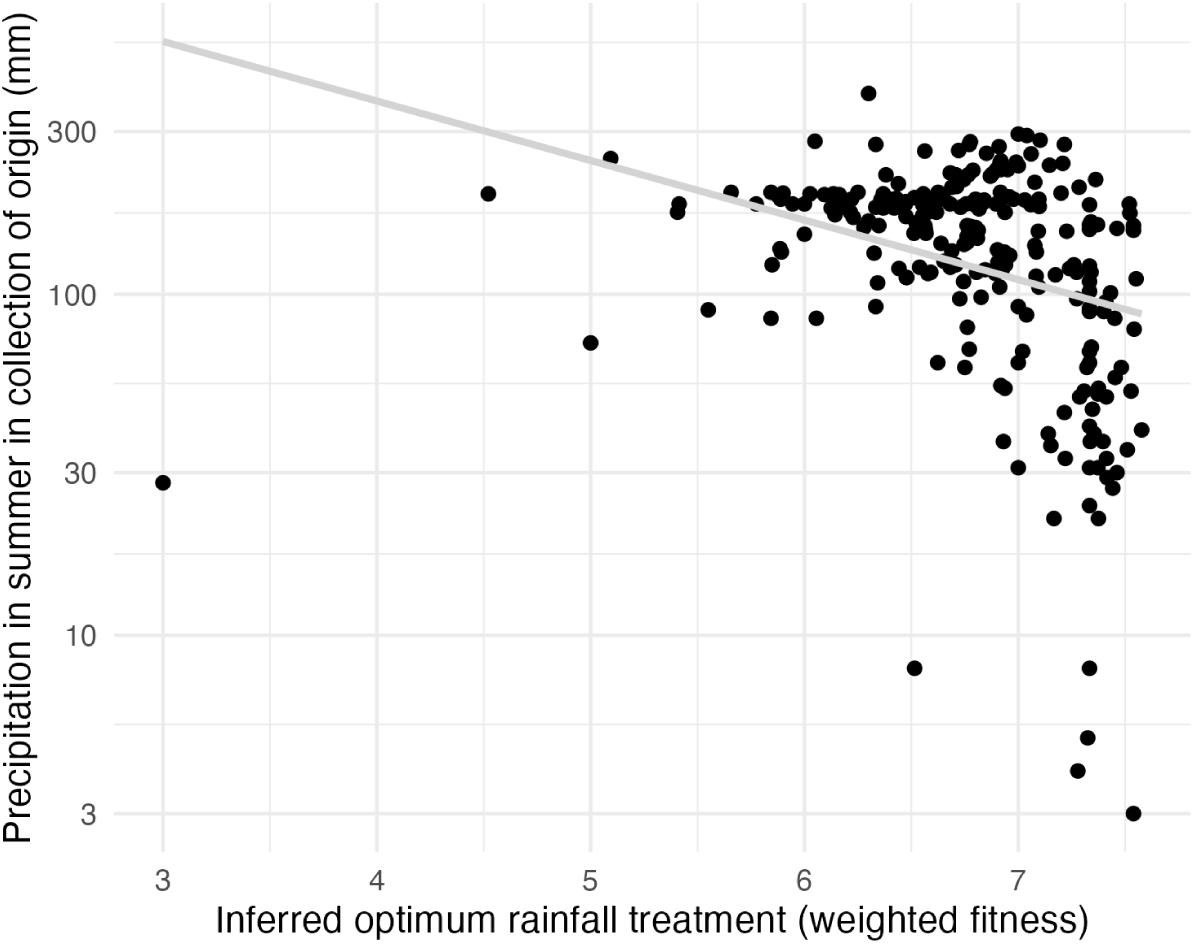
Local adaptation to precipitation in the home environment. Comparison of home site precipitation to the optimum precipitation amount from the field experiment.

**Figure S7.**
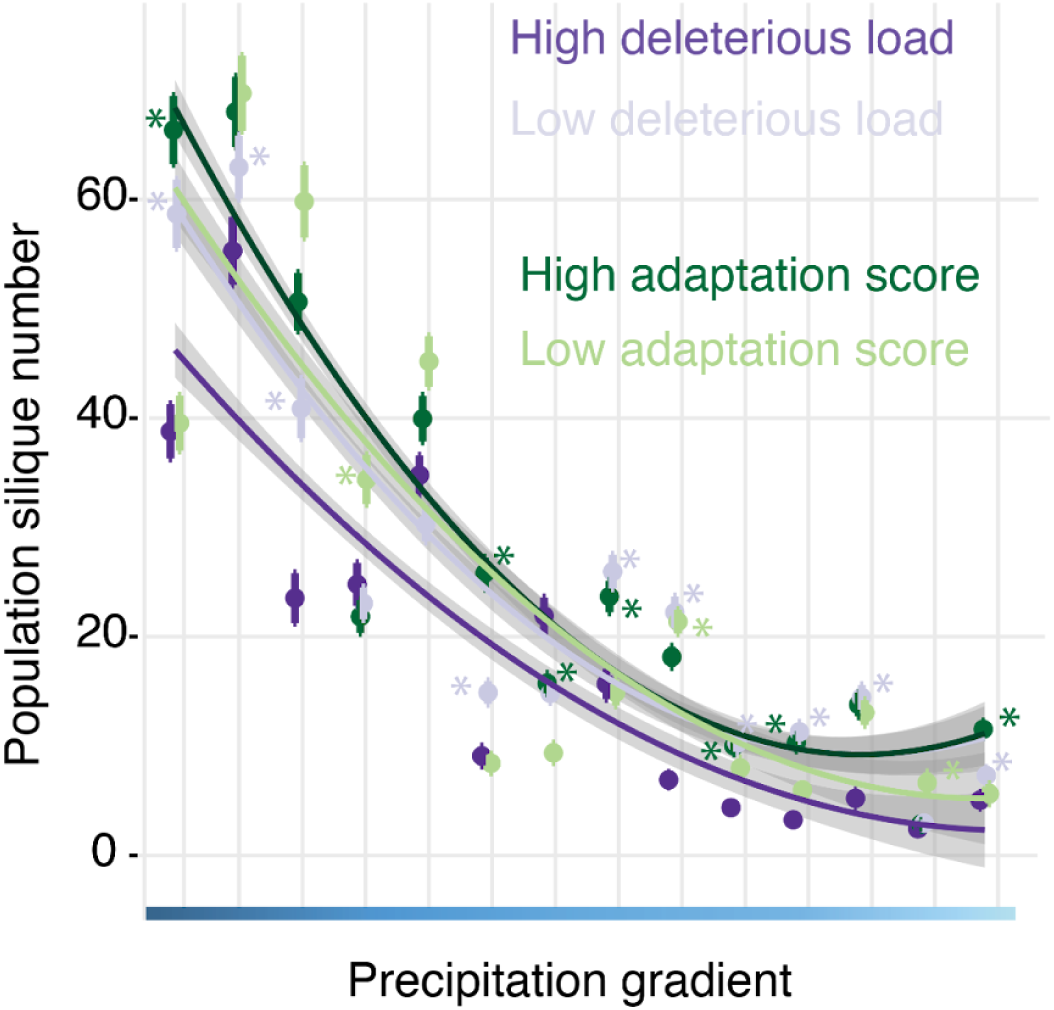
Population-level silique number across 14 precipitation treatments.. Relationship between silique number of tray and precipitation gradient grouped by population. Color indicates the unique population, and paired colors suggest the populations that are paired and that have the Wilcox test conducted on. Asterisks indicate a significant difference (Wilcoxon) in the paired populations.

**Figure S8.**
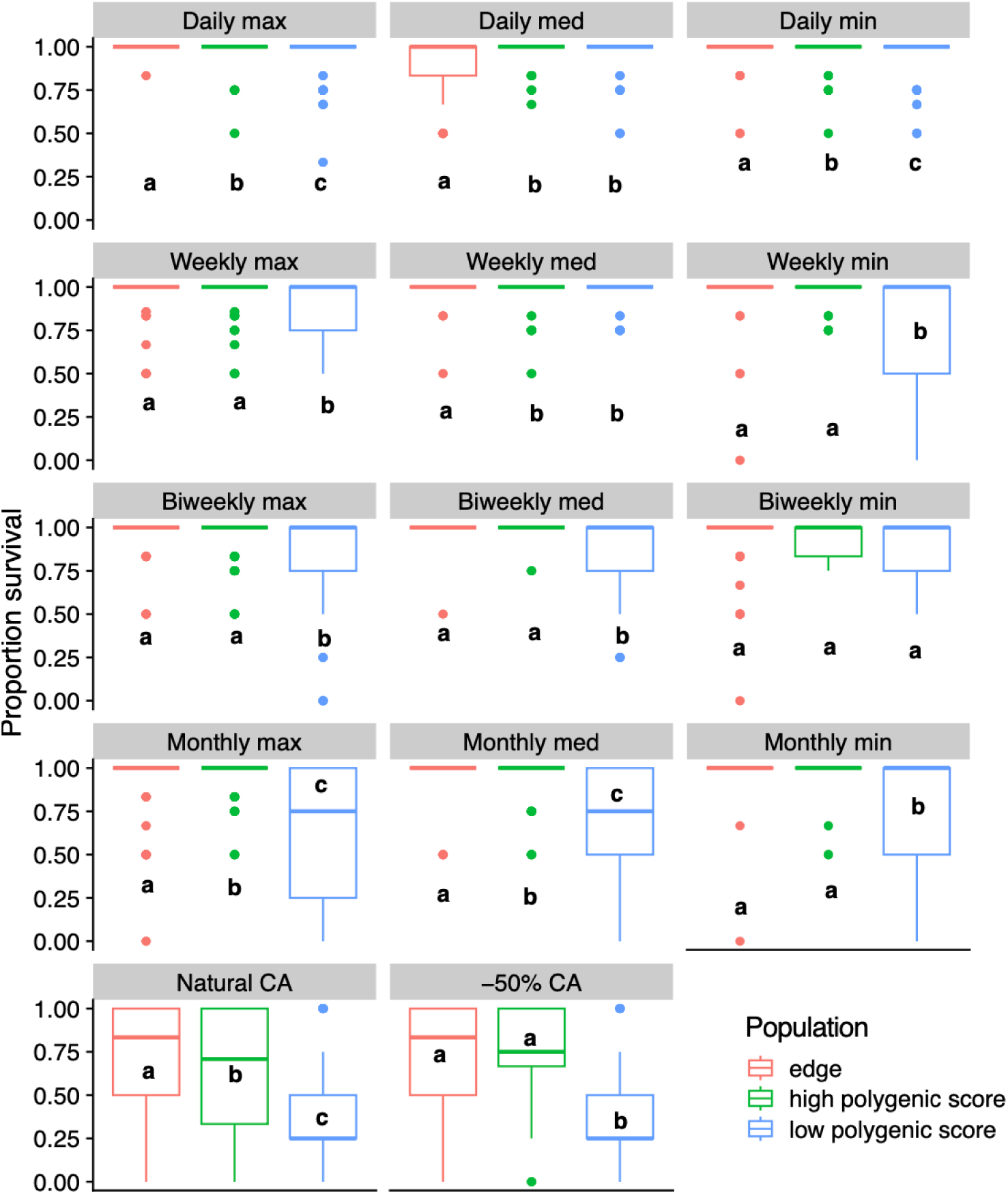
Proportion survival of the high and low adaptation scores populations compared to “edge” populations across treatments. The three boxplots per panel represent the three populations, and a paired Wilcox rank sum test allowed us to determine if the three populations’ fitness were significantly different from each other. Wettest treatment is in the top left corner, and treatments get drier as you go from left to right and down the columns.

**Figure S9.**
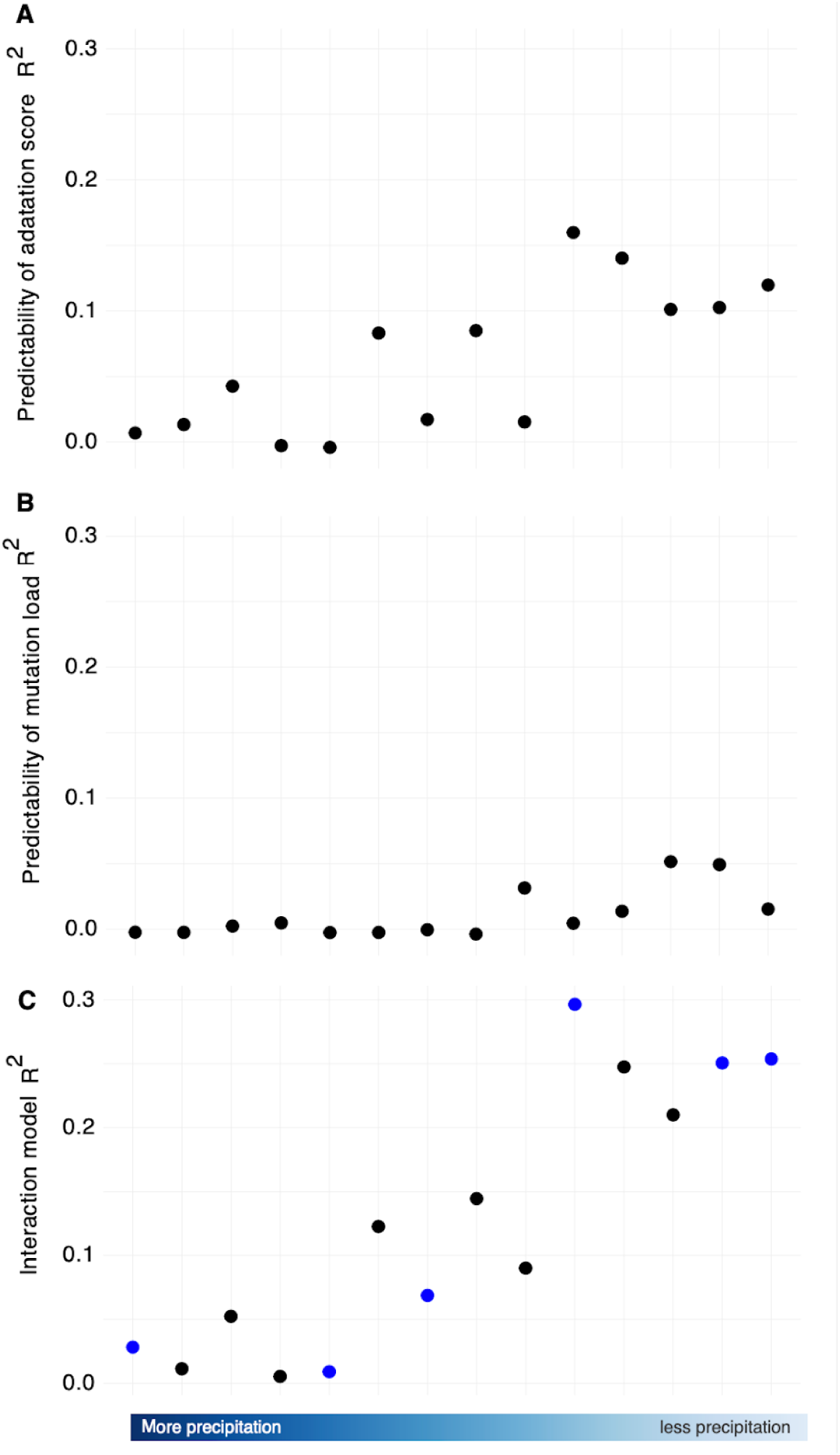
Predictability of genomic measures across treatments. (A) Predictability R^2^ of adaptation score and mutation load (B) across the 14 precipitation treatments. (C) R^2^ from the interaction model of adaptation score and mutation load on fitness within each treatment. Blue dots indicate a significant interaction effect.

**Figure S10.**
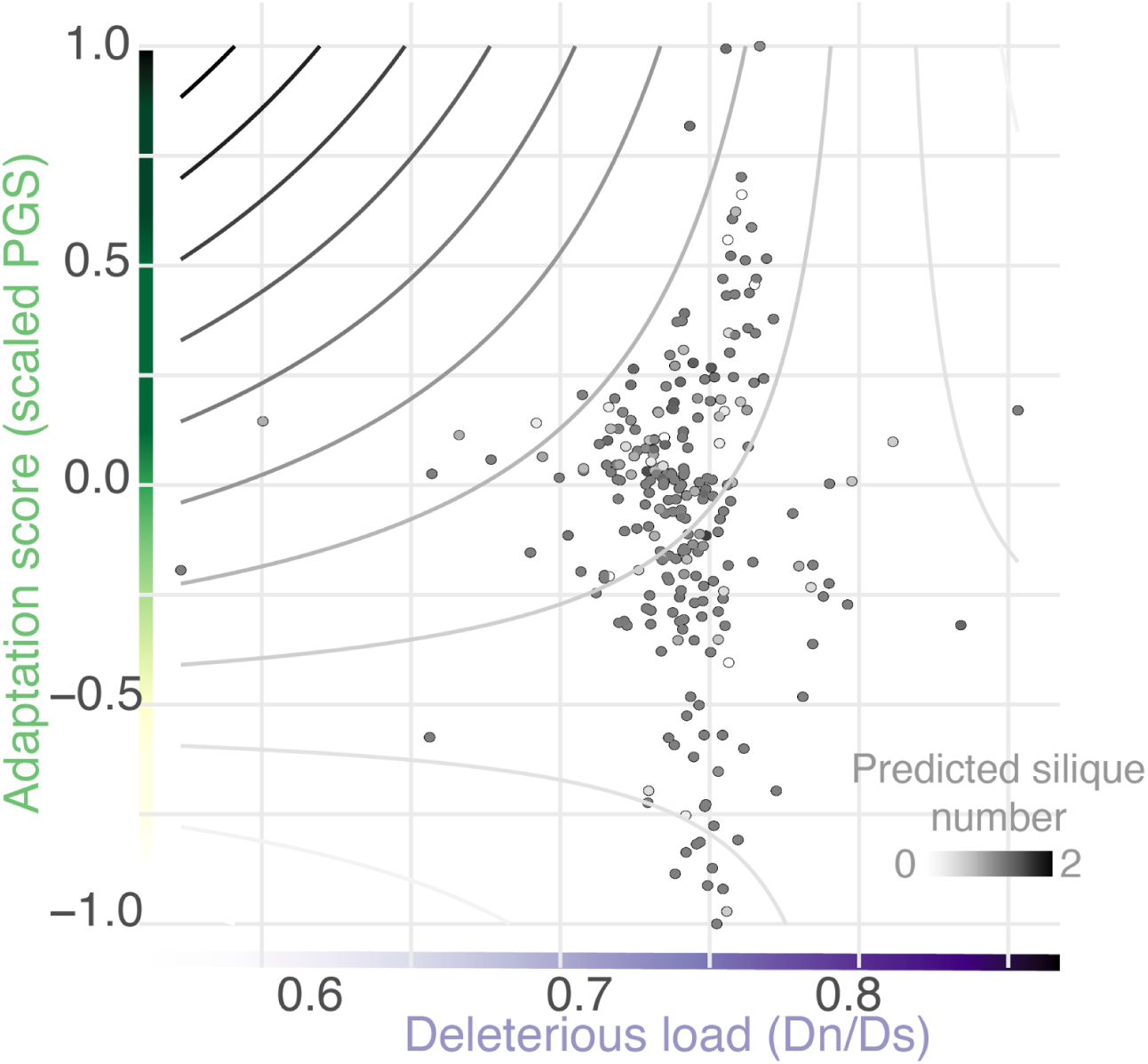
Adaptation score and mutation load colored by silique number in the 50% reduction treatment. Contours highlight the interactive effect of these two genomic measures on fitness.

**Figure S11.**
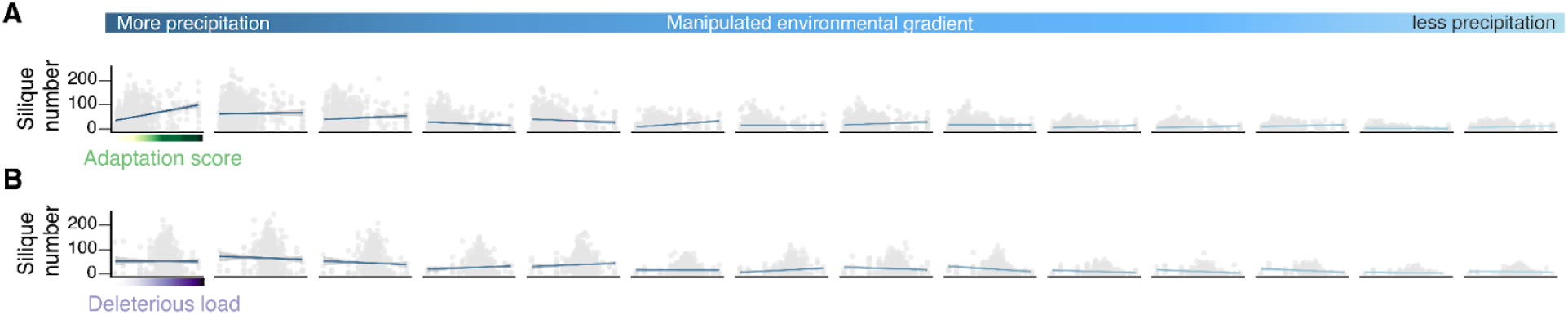
Adaptation score and mutation load relationship with silique number across all treatments. Overall relationship of our two genomic measures (A– adaptation score, B– mutation load) across all accessions by treatment.

**Figure S12.**
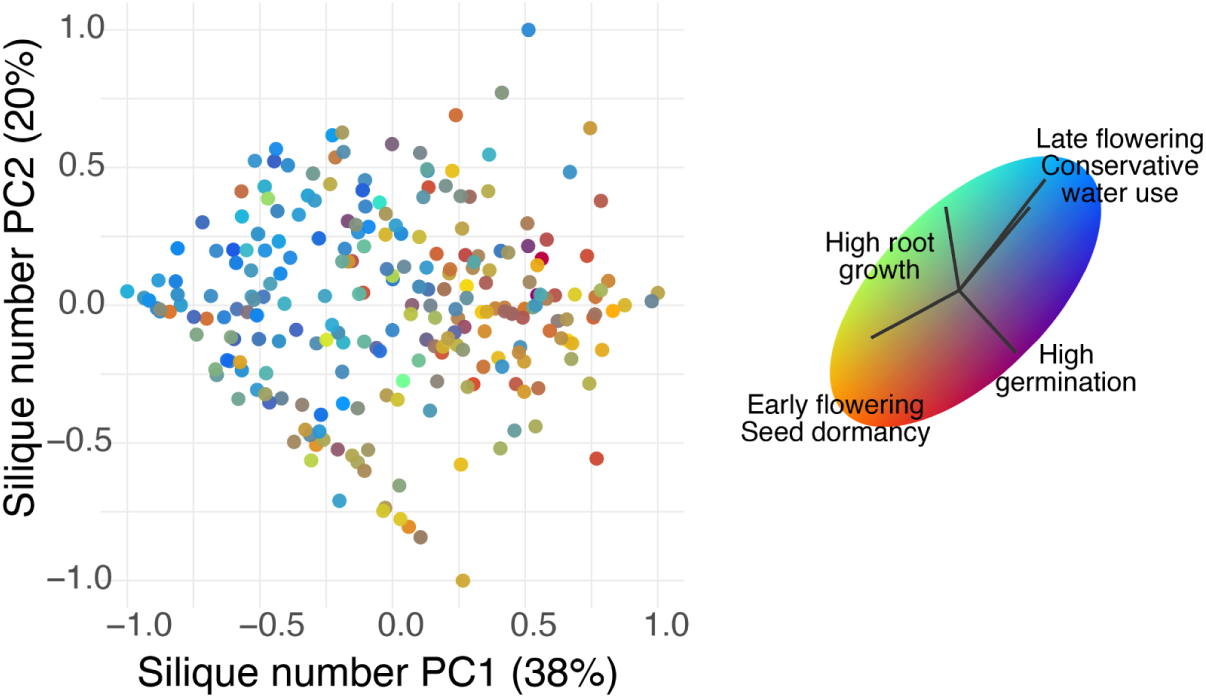
PC1 and PC2 of silique number across treatments colored by phenotype PC. Phenotype PC color range was derived from previous paper (Ruffley et. al 2024), and highlights the fast-flow spectrum of plant growth.

**Figure S13.**
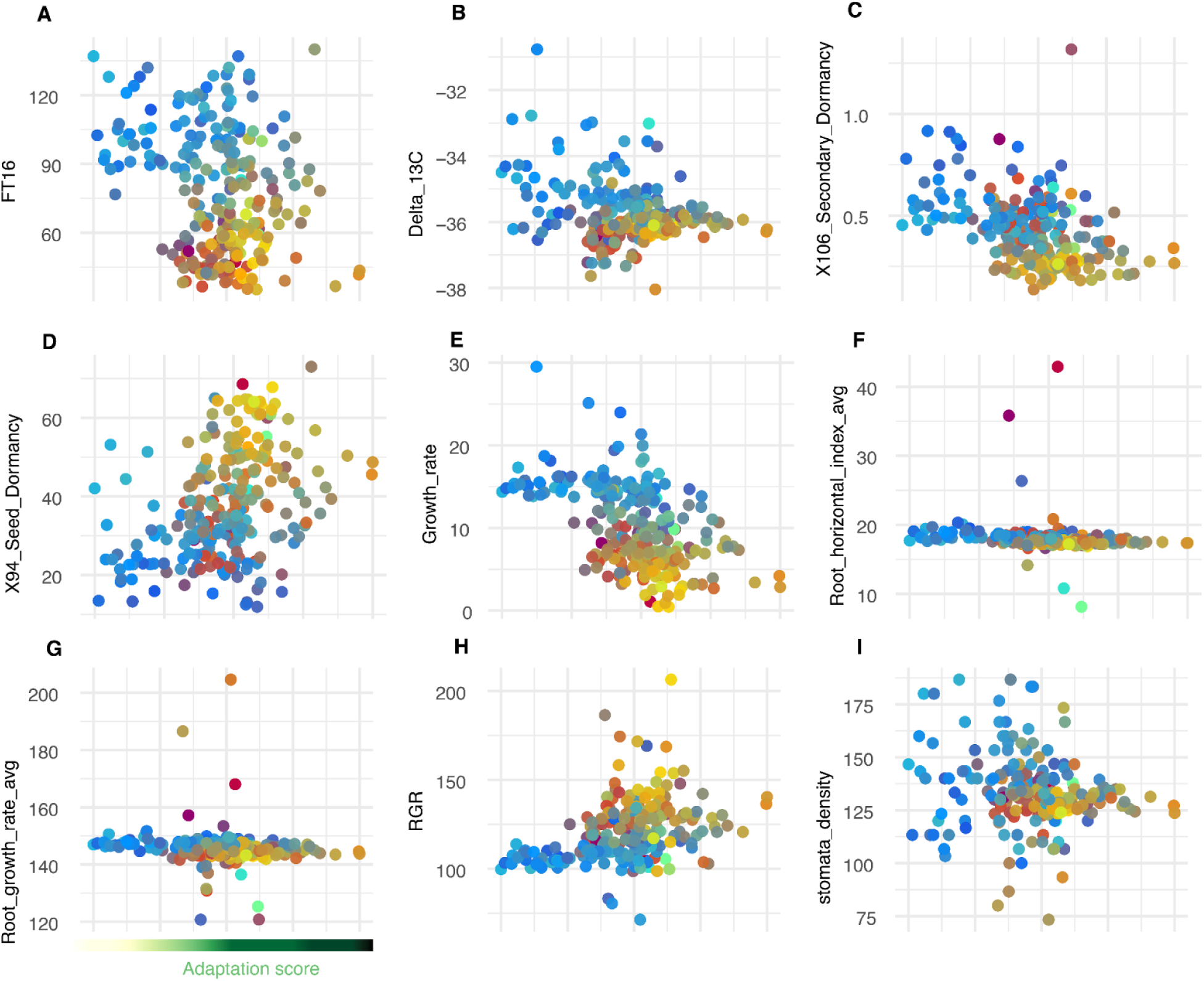
Adaptation score’s relationship with nine get fast-slow phenotypes colored by phenotypic PC1. Phenotypes of interest were selected based on the selection experiment in Ruffley et. al 2024, including (A) Flowering time at 16°C, (B) delta Carbon13, (C) secondary dormancy, (D) seed dormancy, (E) growth rate, (F) average root horizontal index, (G) average root growth rate, (H) relative growth rate, and (I) stomata density. Color of points is from the fast-slow PC-axis in figure S11.

**Figure S14.**
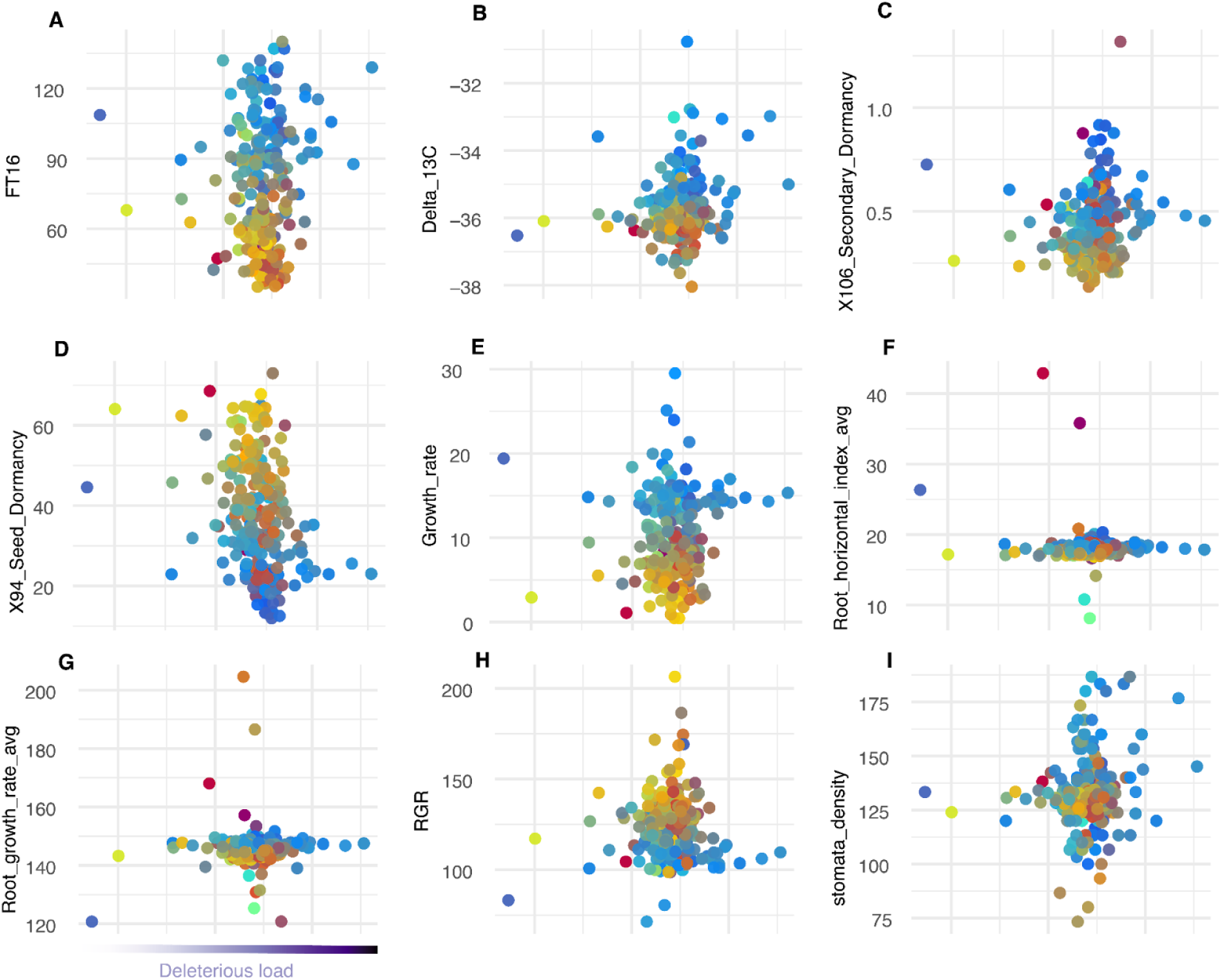
Relationship of mutation load with nine get fast-slow phenotypes colored by phenotypic PC1. Phenotypes of interest were selected based on the selection experiment in ^68^, including (A) Flowering time at 16°C, (B) delta Carbon13, (C) secondary dormancy, (D) seed dormancy, (E) growth rate, (F) average root horizontal index, (G) average root growth rate, (H) relative growth rate, and (I) stomata density. Color of points is from the fast-slow PC-axis in figure S11.

